# Structural and regulatory insights into the glideosome-associated connector from *Toxoplasma gondii*

**DOI:** 10.1101/2023.01.23.525158

**Authors:** Amit Kumar, Oscar Vadas, Nicolas Dos Santos Pacheco, Xu Zhang, Kin Chao, Nicolas Darvill, Helena Ø. Rasmussen, Yingqi Xu, Gloria Lin, Fisentzos A Stylianou, Jan Skov Pedersen, Sarah L. Rouse, Marc L. Morgan, Dominique Soldati-Favre, Steve Matthews

## Abstract

The phylum of Apicomplexa groups intracellular parasites that employ substrate-dependent gliding motility to invade host cells, egress from the infected cells and cross biological barriers. The glideosome associated connector (GAC) is a conserved protein essential to this process. GAC facilitates the association of actin filaments with surface transmembrane adhesins and the efficient transmission of the force generated by myosin translocation of actin to the cell surface substrate. Here, we present the crystal structure of *Toxoplasma gondii* GAC and reveal a unique, supercoiled armadillo repeat region that adopts a closed ring conformation. Characterisation of the membrane binding interface within the C-terminal PH domain as well as an N-terminal fragment necessary for association with F-actin suggest that GAC adopts multiple conformations. A multi-conformational model for assembly of GAC within the glideosome is proposed.

## Introduction

Cellular migration is an essential process that plays important roles in morphogenetic movements, immune cell trafficking, wound healing, and invasion. Interactions between cells and their environment are essential for the transmission of intracellular forces to the extracellular matrix. In multicellular eukaryotes, cell-cell adhesion ensure tissue integrity while providing footholds for the migration of cell within tissues (De Pascalis & Etienne-Manneville, 2017). Cadherins and integrins are major examples of such adhesive molecules that are coupled to the actin cytoskeleton via intracellular bridging components, such as the catenins, vinculin and talin (Bachir *et al*, 2017). Pathogenic organisms also exploit adherent junctions to facilitate movement with respect to host cells and invasion. *Toxoplasma gondii* is an obligate intracellular unicellular parasite and a prominent member of the Apicomplexa phylum (Kim & Weiss, 2004), which also includes Plasmodium, the causative agent of human malaria (Su *et al*, 1995). These parasites share a common set of specialised apical organelles the micronemes and rhoptries, that are critical for invasion (Carruthers & Sibley, 1999). The sequential secretion of both organelles leads to the formation of a moving junction (MJ) formed between the parasite and host cell plasma membranes that participates in active penetration. Apicomplexan parasites also actively egress from infected host cells and migrate across biological barriers. Substrate-dependent, forward parasite propulsion is known as gliding motility and powered by a multiprotein structure referred to as the glideosome. A myosin motor comprising myosin A (MyoA), a class XIV myosin heavy chain, together with glideosome-associated proteins (GAPs) interact with and generate rearward translocation of actin filaments (F-actin) along the parasite (Powell *et al*, 2018).

The glideosome-associated connector protein (GAC) is a central bridging component of the gliding machinery (Jacot *et al*, 2016). This large protein composed of numerous armadillo repeats (ARMs) is highly conserved throughout the Apicomplexa phylum and links F-actin to the TRAP/MIC family of surface adhesins in the plasma membrane, which target host cell ligands and mediate adherent anchor points. GAC translocates dynamically with the MJ from the parasite apical to the basal pole during gliding motility, host cell egress and invasion. Rearward translocation of adhesins anchored to both the parasite plasma membrane and the host membrane by the inner-membrane-associated glideosome generates parasite forward movement (Carruthers & Tomley, 2008; Frénal *et al*, 2017a).

The initial apical location of GAC also depends on the activity of an apical lysine methyltransferase (AKMT), through a yet unknown mechanism (Jacot *et al*., 2016). Recently ultrastructure expansion microscopy allowed to position GAC and formin 1 (FRM1) to the preconoidal rings (PCRs) (Dos Santos Pacheco *et al*, 2022). Importantly, FRM1 is the only and essential nucleator of actin polymerization to drive conoid extrusion and parasite motility and invasion (Tosetti *et al*, 2019). As part of the conoid the PCRs serve as platform for the assembly of the glideosome. Membrane-association of GAC relies on its capacity to bind phosphatidic acid (PA), an essential lipid mediator for microneme secretion which assists in the correct engagement of GAC (Bullen *et al*, 2016). Despite significant efforts, high-resolution structural insight into GAC and its multiple interactions has not been available since its discovery. An initial SAXS model predicted an elongated club-shaped conformation with the C-terminal PH domain lying at the extremity of the structure.

Here, we describe combined X-ray crystallography, nuclear magnetic resonance (NMR) and small angle X-ray scattering (SAXS), Hydrogen/Deuterium exchange coupled to Mass Spectrometry (HDX-MS) and course-grained molecular dynamics (CG-MD) analyses to illuminate the structure and conformations adopted by full-length GAC from *T. gondii*. Structure validation by biochemical and parasite assays provides insights into membrane and actin binding ultimately proposing a model for assembly within the glideosome.

## Results

### The structure of full-length TgGAC

Well-diffracting and reproducible crystals were obtained at pH 5 for native full-length TgGAC (residues 1-2639) (Kumar *et al*, 2022). Selenomethionine-substituted crystals were also obtained that produced sufficient anomalous signal for phase determination. The structure was solved by multiple-wavelength anomalous dispersion (MAD) to 2.7 Å resolution with an Rfree of 26% (Table 1). The electron density for residues 7-2504 was of sufficient quality to facilitate modelling for these residues (Figure S1a). The remaining electron density showed evidence peptide backbone but could not be confidently modelled and refined, indicating a degree of conformational flexibility for residues 2505-2639.

**Table 1.** Data collection, phasing and refinement statistics

|  | TgGAC |
| --- | --- |
| Space group | P 21 21 21 |
| Cell dimensions |  |
| *a*, *b*, *c* (Å) | 119.078 123.605 221.508 |
| $\alpha$, $\beta$, $\gamma$ (°) | 90, 90, 90 |
| Wavelength | 0.97625 |
| Resolution range (Å) | 55.95 - 2.675 (2.771 - 2.675) |
| $R_{pim}$ | 0.04101 (0.3725) |
| $I / \sigma I$ | 14.10 (0.71) |
| Completeness (%) | 99.90 (99.42) |
| Wilson B-factor | 84.96 |
| R-merge | 0.1225 (1.113) |
| R-meas | 0.1294 (1.175) |
| CC/12 | 0.993 (0.681) |
| CC* | 0.998 (0.9) |
| No. reflections for refinement | 92809 (9144) |
| No. reflections for $R_{free}$ | 4517 (395) |
| $R_{work}$ | 0.2237 (0.4037) |
| $R_{free}$ | 0.2694 (0.4463) |
| CC (*work*) | 0.897 (0.564) |
| CC (*free*) | 0.917 (0.447) |
| No. non-hydrogen atoms | 18441 |
| Macromolecules | 18346 |
| Solvent | 95 |
| Ramachandran favoured /<br>allowed/ outliers (%) | 92.3/ 6.0/1.7 |
| Average B-factor | 83.47 |
| Macromolecules | 83.53 |
| Solvent | 71.30 |
| Rms Bond lengths (Å) | 0.013 |
| Rms Bond angles (°) | 1.3 |
| Clashscore | 17.98 |

\*Data from a single crystal were used to solve the structure.
\*Values in parentheses are for highest-resolution shell.

To gain insight into the structure of this C-terminal region, we initiated a solution NMR spectroscopy approach with a construct encompassing residues 2505-2639 (TgGAC_2505-2639_). NMR spectra confirmed an independently folded domain and 80% of the backbone resonances could be confidently assigned (Figure S1b). Several amide peaks were broadened or absent from the spectra due to conformational exchange in some loop regions. We next generated a structural model for the TgGAC_2505-2639_ domain using AlphaFold2 and validated these with available NMR chemical shifts and nuclear Overhauser effects (NOE) data. Excellent agreement was observed between NOEs predicted from the Alphafold2 structure and those observed in NOESY spectra of TgGAC_2505-2639_ (Figures S1c–S1e).

The structure of TgGAC_2505-2639_ adopts a PH-like domain fold comprising a 7-stranded β-barrel with 3 a-helices. The presence of an extended N-terminal-helix is reminiscent of the PH domains from TgAPH and TgISP (Darvill *et al*, 2018; Tonkin *et al*, 2014) and appears to be a common feature among apicomplexan PH domains. To complete a model for full-length TgGAC, the validated structure of TgGAC_2505-2639_ (TgGAC_PH_) was positioned in the available electron density map for this region and a linker modelled with MODELLER (Fiser *et al*, 2000).

The full-length TgGAC structure comprises 169 a-helices (Figure S2) arranged in 53 consecutive helical bundles that resemble Armadillo (ARM) and HEAT-like repeats (AHRs) (Kippert & Gerloff, 2009) linked to the mixed α/β C-terminal PH domain (Figure 1a). The architecture can be divided into three major regions (RI, RII and RIII). RI (residues 1 to 1665) comprises 37 consecutive AHRs of approximately 40 residues long that are supercoiled into a 3-layer pyramid structure (Figure 2a). The final ARM of region I (AHR 37) is extended with a 6 amino acid linker (L1) to start a second large AHR region (RII). RII comprise 16 canonical armadillo repeats (AHR38-53) and forms a superhelical arch that contacts the base of the RI pyramid helical regions. This interface encompasses a surface area of 1550Å^2^, with AHR1 and AHR17-18 from RI forming one side while AHR48-50 from RII form the other (Figure 1c). The interface comprises numerous hydrogen bonds and key salt bridges (K2254-D11, E2436-H24, D2330-R629, D2346-K672, D2197-R716; Figure 1d). AHR50 also has a large helixloop-helix insertion (residues 2278 to 2337) between the 2^nd^ and 3^rd^ helices which creates a prominent protrusion that also stabilises the interface (Figure 1b & 1c). A basic AHR53 completes region (II) and an ordered linker from 2489 to 2510 (L2) extends to the adjacent C-terminal PH domain which comprises region III (Figure 1a & 1b).

**Figure 1.**
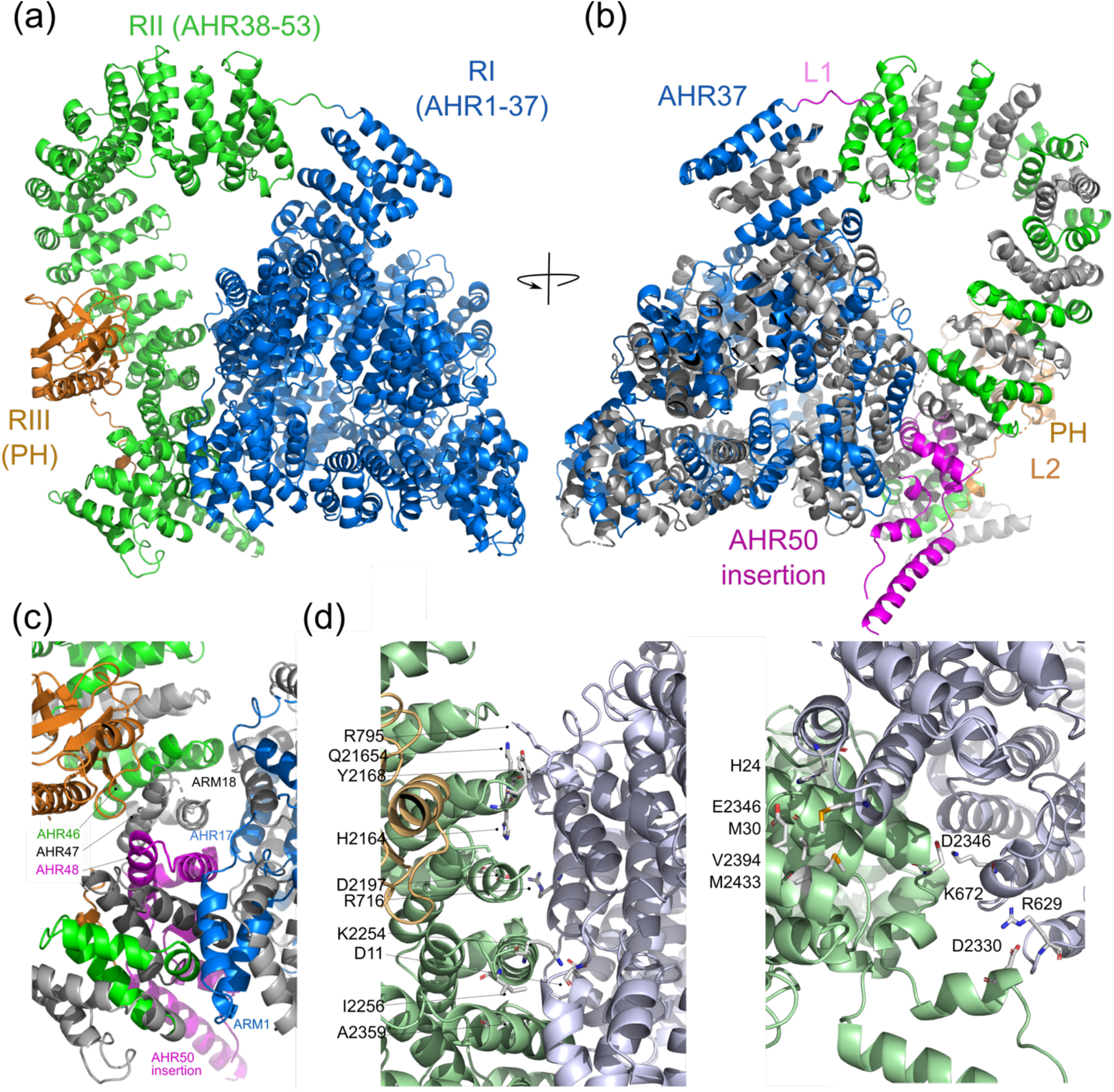
Cartoon representation of the TgGAC structure. (a) Overall architecture. Region I (R1: residues 1 to 1665) comprising the first 37 consecutive armadillo (ARM)/HEAT-like repeats (AHRs) is supercoiled into a 3-layer pyramid structure (blue). AHR region II comprising 16 ARM repeats (RII: AHR38-53 residues 1670-2489) forms the superhelical arch (green). The C-terminal PH domain encompassing 2511-2639 of RIII is shown in orange. (b) 180° rotation of the orientation shown in (a) with even numbered AHRs shown in grey and odd number shown in blue for RI and green for RII. The R1-RII linker and AHR50 which has a helix-loop-helix insertion are shown in magenta. (c) The N/C interface between RI and RII showing key interacting AHRs. (d) Key residues specific interactions across the N/C interface. Cartoon representations coloured in light blue for RI and light green for RII.

**Figure 2.**
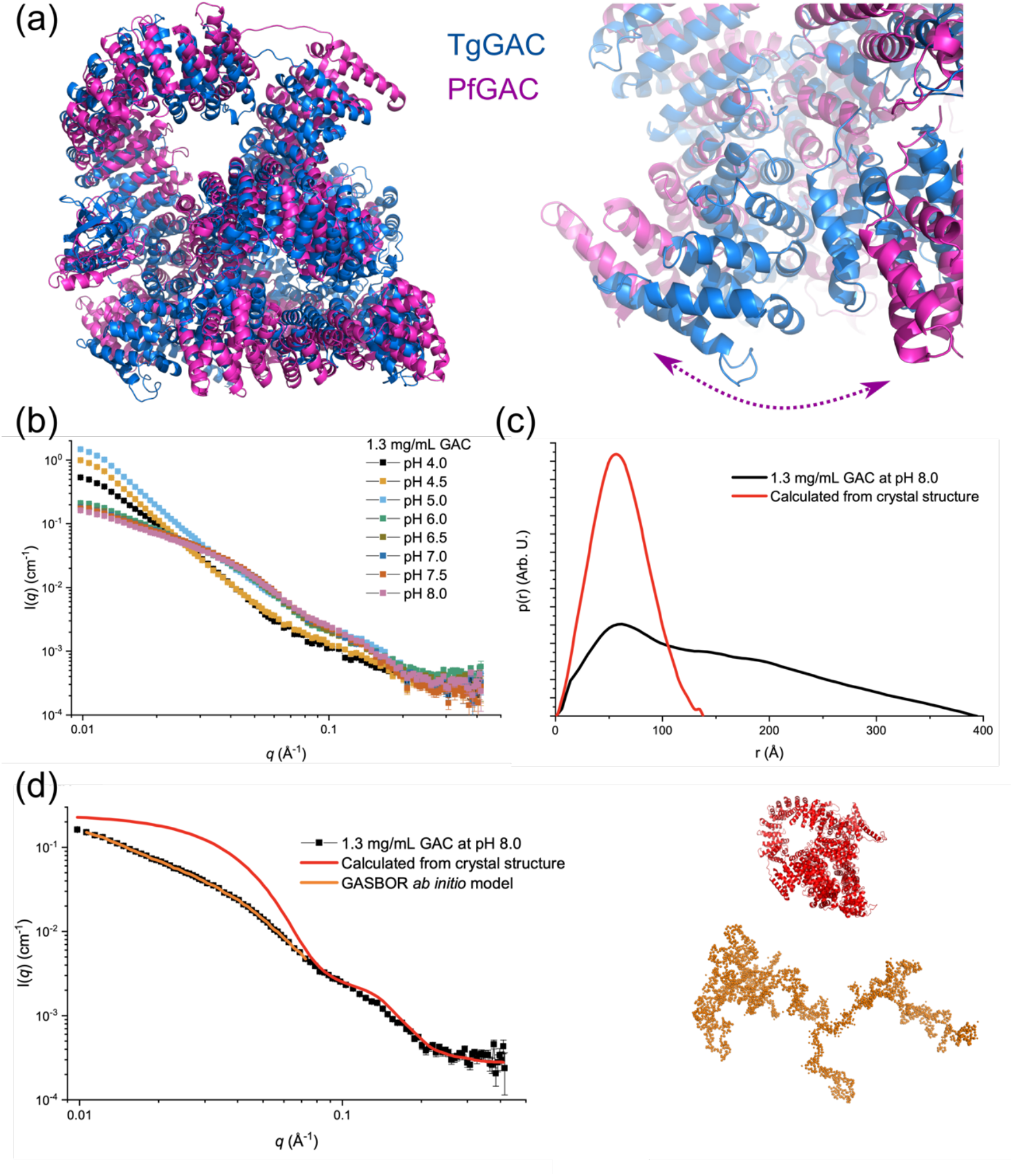
GAC adopt both open and closed structures. (a) Superimposition of the crystal structure of TgGAC with that generates from Alphafold for PfGAC shows the closed structure is conserved at the sequence level (left). Interface is highly similar but partially opened in the predicted structure (right). (b) SAXS data for GAC at pH 4.0 – 8.0. (c) p(*r*) function for GAC SAXS data and calculated from crystal structure. (d) Modelling of GAC solution structure at pH 8.0. The red curve is the calculated theoretical scattering curve for the crystal structure shown in red. GASBOR ab initio model and the fits are shown in orange.

### GAC adopts multiple extended conformations in solution

The closed conformation observed in the TgGAC crystal structure deviates significantly from the extended club-shaped structure that was proposed from previous SAXS analyses (Jacot *et al*., 2016), which raises concern that it could be an artefact of crystallisation. To shed light on this, we examined an AlphaFold2 structure predicted for full-length PfGAC (Jumper *et al*, 2021; Varadi *et al*, 2022).

Remarkably, the overall architecture of the PfGAC predicted structure is similar to the one observed for TgGAC, as it is characterised by a series of tandem AHR repeats that are supercoiled into a large ring, closed by specific interactions between N- and C-terminal AHRs. While the interface in PfGAC is partially separated (Figure 2a), the interacting regions overlap with those identified experimentally for TgGAC.

We next performed a SAXS analysis over a range of concentrations and solution conditions to determine whether an open structure is indeed present in solution. GAC was measured at a concentration of 1.3 mg/mL as a function of pH from 4.0 to 8.0. The largest difference is observed as an increase in intensity at low scattering vector moduli, *q* (Figure 2b), where the intensity is proportional to the mass of the protein/complexes, suggesting that oligomerization is occurring and is most pronounced at pH 5.0.

To explore the monomeric structure of TgGAC in solution at pH 8.0, the mass and size were determined both by a Guinier fit (Figure S2) and using an indirect Fourier transform (IFT) routine for calculating the pair distance distribution function, *p*(*r*) (Figure 2c). Masses of 264 kDa and 286 kDa were obtained, which are close to the expected value of 286 kDa for a TgGAC monomer. Radius of gyration, *R*_g_, values of, respectively, 105 Å and 122 Å were obtained, which are much larger than the *R*_g_ of 48 Å calculated from the crystal structure of TgGAC. This discrepancy in size is also clearly seen when comparing the *p*(*r*) function calculated from the SAXS data or the crystal structure (Figure 2c). The theoretical scattering curve can be calculated from the PDB structure and compared to the measured data (Figure 2d). It shows that at medium to large *q*, the structures are relatively similar, indicating that they contain similar structural components on shorter length scales, but at medium to low *q*, there is a large difference showing that the overall size and shape are very different. To explore this further, an *ab initio* model was build using the ATSAS program GASBOR (Svergun *et al*, 2001), which is clearly highly extended and fits the low *q* part of the experimental data (Figure 2d). Lastly, a dimensionless Kratky plot illustrates that the structure in pH 8.0 solution is much more flexible than would be predicted from the crystal structure, but is not completely unfolded (Figure S3), Overall, these data show that even though TgGAC is primarily a monomer in solution at pH 8.0, it forms a oligomerized and more extended structure than seen in the crystal structure whilst maintaining key secondary-structural elements. Collectively, it can be concluded that a closed structure is likely to be an important stable functional state as the N/C-interface is encoded within the sequence conservation. The extended, flexible structure observed in solution may play a role is a facilitating GAC’s recruitment of its binding partners.

### Mapping the PA-binding interface for the C-terminal PH domain of GAC

An extended patch of conserved basic residues is formed on one face of the PH domain in GAC which is reminiscent of the charge distribution on TgAPH (Darvill *et al*., 2018) suggesting a similar role in phospholipid binding. Using phospholipid strip assays, it had been previously demonstrated that both PfGAC and TgGAC bind specifically to phosphatidic acid (PA) (Jacot *et al*., 2016). Building upon our earlier work on the PA binding protein TgAPH (Darvill *et al*., 2018), we employed coarse-grained molecular dynamics (CG-MD) simulations to characterise the binding of TgGAC to PA within a membrane environment. In all three simulation repeats, GAC bound to the membrane within 10 μs (Figure S3). Analysis of protein-lipid contacts highlight two key membrane-bound regions, which localised to the PH domain and an adjacent basic protrusion from AHR53 (Figure 3).

**Figure 3.**
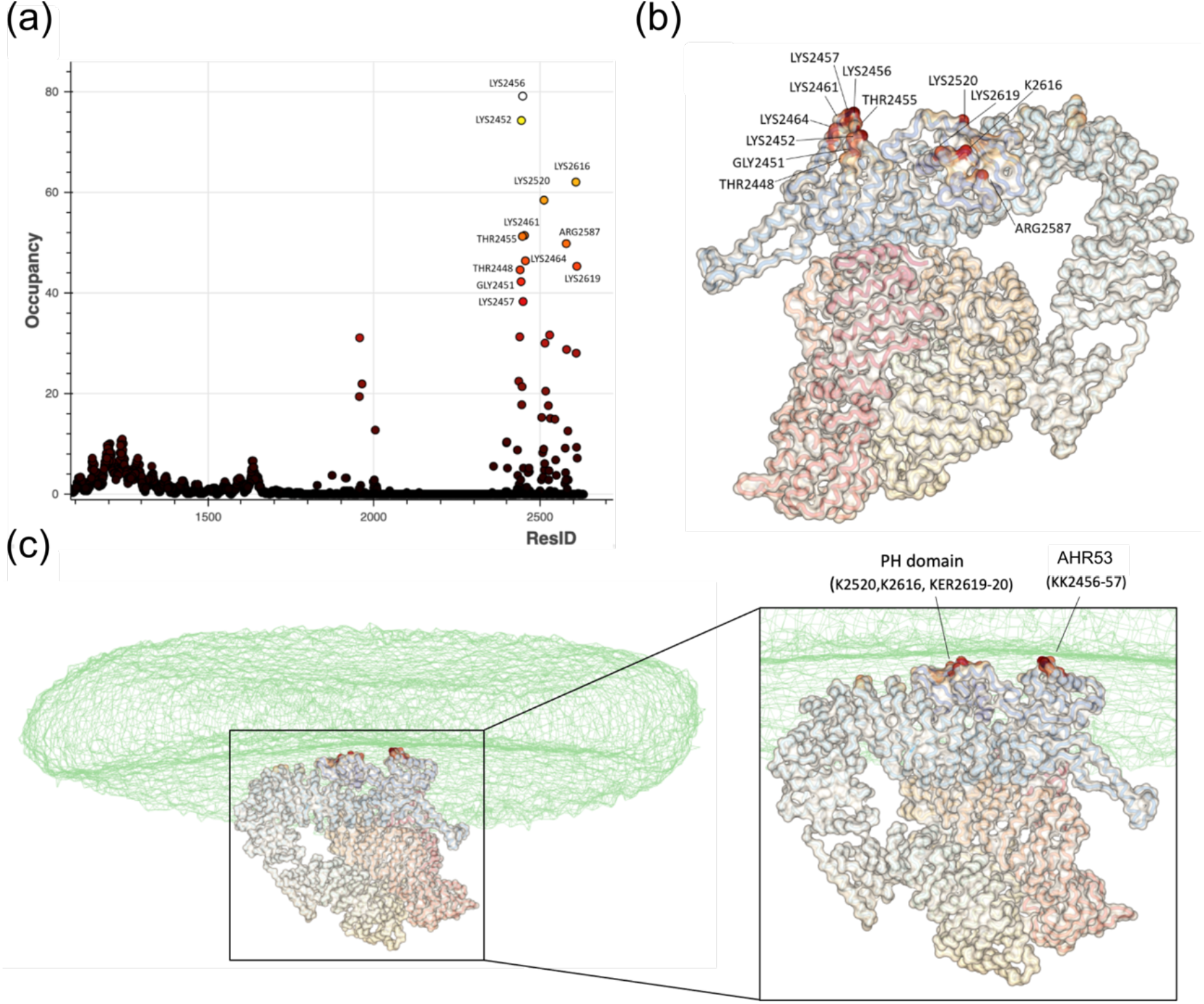
Simulation of TgGAC binding to PA-containing membranes. (a) Analysis of residues that interact with POPA as a percentage of contact during the simulation time. Analysis was performed for the final 5 us of the simulation. (b) Key residues from mapped onto the TgGAC structure. (c) As in (b) in the contact of the membrane which is displayed as a transparent surface.

To complement out NMR studies on TgGAC_PH_ and enable a more comprehensive map of the PA-binding interface, we also explored the equivalent region from *Plasmodium falciparum* (53% identity over PfGAC_2471-2605_) with NMR mapping studies. The quality of the NMR spectra was sufficient to confidently assigned over 95% of the backbone residues and validate the Alphafold models (Figure S4a). Exploiting the near complete NMR assignment of PfGAC_PH_ and its high sequence conservation, we employed NMR titrations to quantify and map the PA binding site. First, 1D NMR binding assays were performed with large PA-enriched MSP1E1 nanodiscs to measure the affinity of the interaction. Titration with MSP1E1 nanodiscs composed of POPC and POPA (50%:50% POPC:POPA) caused a significant loss in PfGAC_PH_ NMR signals (Figure 4a) indicating binding to the ‘NMR invisible’ nanodiscs, whilst almost no loss in signal is observed upon titration with LUVs composed solely of POPC. Binding curves were generated from integration of the NMR signals and (Figure 4b) apparent dissociation constant K_d_s calculated (Figure 4c). The K_d_ for PfGAC_PH_ binding LUVs composed of 50% POPA is calculated to be 60 ± 3μM.

**Figure 4.**
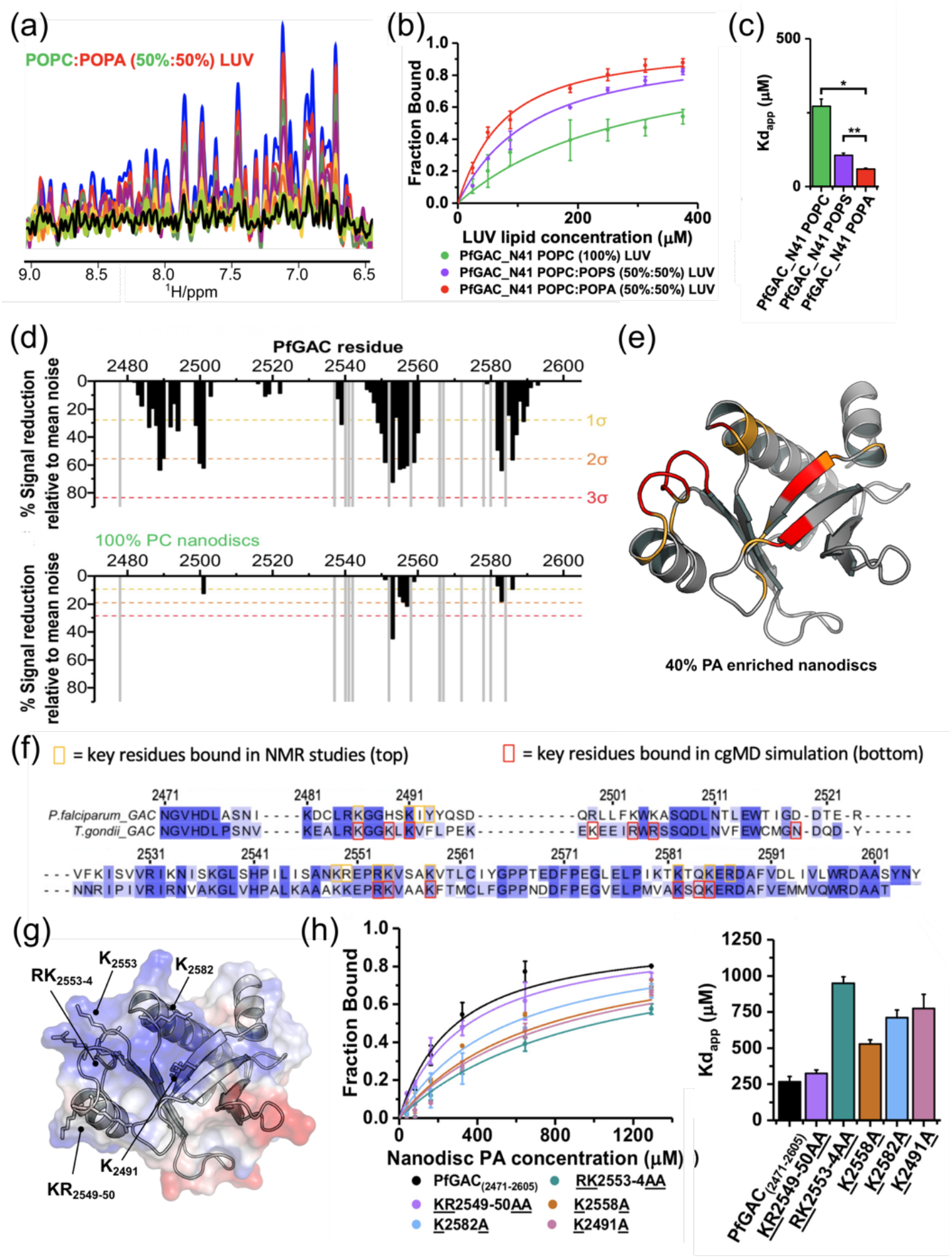
NMR-based binding assays for PfGAC_2471-2605_ (PfGACPH) to PA-enriched unilamellar liposomes. (a) 1D 1H-NMR spectrum (9.4 to 6.5ppm) of PfGACPH upon titration with increasing concentrations of LUVs composed of POPC:POPA (50%:50%) LUV molar ratios: blue, free PfGACPH in solution; red 1:2; green 1:4; purple 1:7; yellow 1:15; orange 1:20; lime 1:25; black 1:30. (b) Binding curves generated from spectral integration and expressed as the fraction of bound protein for variable LUV compositions (POPC (100%) green, POPC:POPS (50%:50%) purple, or POPC:POPA (50%:50%) red). (c) Apparent dissociation constants (K_dapp_) for binding LUVs were calculated from fitting binding curves. (d) Plot of PfGACPH PREs with PA-enriched PA-enriched MSP1D1H4-5 nanodiscs top (POPC:POPA:PE-DTPA-Gd3+ 46%:40%:14%) and bottom (POPC:PE-DTPA-Gd3+ 86%:14%), against sequence number. Dashed lines represent 1 (yellow), 2 (orange), or 3 (red) standard deviations from the mean noise (baseline). (e) PA PREs mapped onto the structure of PfGAC_PH_, residues and coloured if greater than 2 (orange), or 3 (red) (e) Comparison of contact resides from MD and NMR mapped on to the sequence alignment between PfGAC_PH_ and TgGAC_PH_ (g) Electrostatic surface representation of PfGACPH revealing an extensive surface patch of positive charge surface charge with key mutated residues labelled. (h) Binding curves of PfGAC_PH_ mutants generated from 1D NMR titration with calculated K_dapp^S^_

We next measured paramagnetic relaxation enhancements (PREs) using small PA-enriched MSP1D1_δH4-5_ nanodiscs. PA-enriched MSP1D1_ΔH4-5_ nanodiscs (40%:60%, POPA:POPC) or nanodiscs containing no PA (100% POPC), doped with (paramagnetic nanodiscs) or without (diamagnetic nanodiscs) PEDTPA-Gd^3+^ paramagnetic lipid, were generated. 2D ^1^H-^15^N HSQC spectra were recorded for ^15^N-labelled PfGAC_PH_ in the presence of paramagnetic or diamagnetic nanodiscs. Paramagnetic induced relaxation enhancements of membrane-interacting regions were measured from reductions in signal intensities. Numerous PREs were observed for PfGAC_PH_ upon the addition of PA-enriched paramagnetic nanodiscs (Figure 4d - top) but not with 100% POPC nanodiscs (Figure 4d - bottom).

PREs mapped onto the structure of PfGAC_PH_ reveal a contiguous surface formed from residues located within the β1-strand, bordering the β1-β2 loop, within the β5-β6 loop and the loop region between β7 strand and C-terminal α-helix (Figure 4e). A comparative analysis of the PH binding residues compared to simulation with fulllength TgGAC indicated that the PH domain alone binds with the same interface in the context of the whole GAC protein (Figure 4f). NMR titrations performed with TgGAC_PH_ and PA-enriched paramagnetic nanodiscs confirmed an identical PA-binding surface to PfGAC (Figure S4b). Collectively, these results suggest that the solvent exposed positively charge surface in the GAC PH domain represents a PA-specific membrane interacting interface (Figure 4g). Conserved basic residue sidechains are likely to coordinate PA lipid phosphate head groups. A series of alanine substitution mutants were then generated in PfGAC_PH_ for residues identified from NMR mapping. These mutants were tested using the 1D NMR binding assay to quantitatively assess the consequence of mutations on PA-binding residues (Figure 4h).

Compared to WT (K_d_ of 270 ± 40 μM), mutation of basic residues reduced the affinity for PA-enriched MSP1E1 nanodiscs, indicating that these residues are important to binding PA within a membrane environment (Figure 4h). The reduction in affinity correlates with position of basic residues relative to the centre of the solvent exposed positive charges, i.e. the largest effect is observed for the RK_2553-54_AA mutant (K_d_ of 950 ± 50 μM) compared to the KR2549-50AA mutant (K_d_ of 330 ± 20 μM). The same trend is observed for K2491A (K_d_ of 770 ± 100 μM) and K2582A (K_d_ of 710 ± 50 μM), which are more centrally located than K2558A (K_d_ of 550 ± 50 μM).

To establish whether the PA binding interface identified through NMR analyses on PfGAC_PH_ mutants display a similar deficiency in full-length TgGAC, we exploited a liposome binding assay in which bound protein was quantified by sedimentation and SDS-PAGE. We first tested this assay for full-length TgGAC, TgGAC_PH_ and the known PA sensor TgAPH (Darvill *et al*., 2018) in presence of liposomes with no PA and liposomes containing 50% PA. All three proteins were bound specifically to PA-enriched liposomes with most of the protein present in the pellet after ultracentrifugation, whereas in the absence of PA the proteins were found in the supernatant (Figure 5a). Three mutants were chosen based on the NMR data generated with both full-length TgGAC and isolated PH domain constructs. The first mutant focusses on the major positive charged patch that exhibited the largest effect in NMR assays (namely RK_2553-54_AA), the second is a triple mutation in the downstream basic region (KER_2585-2587_AAA) and the third combines both (RK_2553-54_AA/KER_2585-2587_AAA). All mutant proteins tested experience a significant reduction in binding to PA enriched liposomes (figure 5b and 5c), with the most dramatic effect observed for the quintuple mutant RK_2553-54_AA/KER_2585-2587_AAA for both GAC PH domain and fulllength.

**Figure 5.**
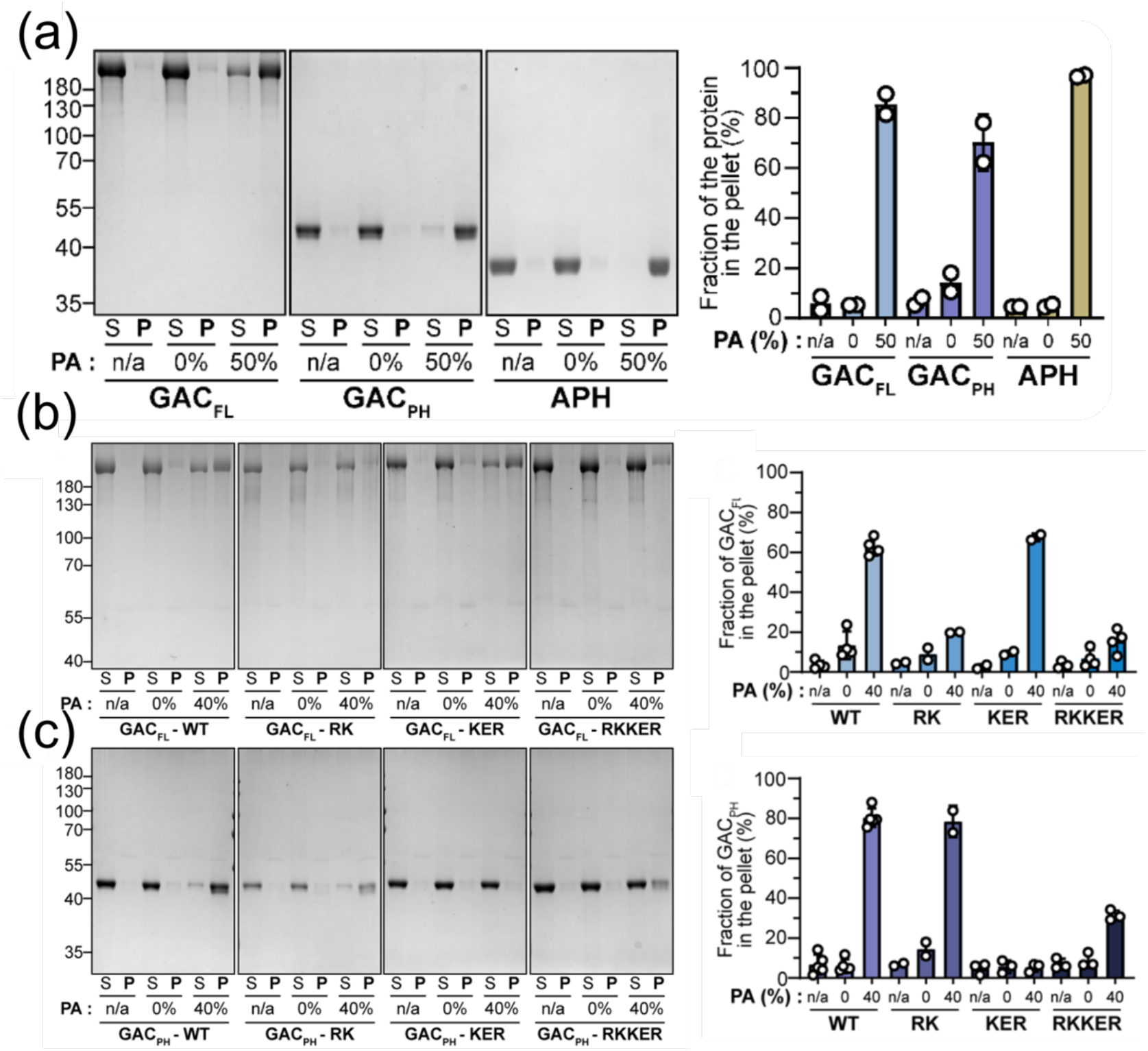
Liposome binding assays of PA binding by GAC and PH domain mutants *in vitro*. (a) Liposome binding assay with the three proteins of interest. A representative gel stained by Coomassie blue (left). Quantification of the pellet fraction measured by band densitometry (n=2). n/a= no liposome. S=supernatant fraction after ultracentrifugation. P = pellet fraction after centrifugation (right). (b) Representative gel stained by Coomassie blue from liposome binding assays with full-length TgGAC_FL_ and the three mutated versions. n/a = no liposome. S = supernatant fraction after ultracentrifugation. P = pellet fraction after centrifugation. RK = RK/AA mutations. KER = KER/AAA mutations. RKKER = RK/AA + KER/AAA mutations. (c) Representative gel stained by Coomassie blue from liposome binding assays with TgGAC_PH_ and the three mutated versions. (d) Quantification of the GAC_FL_ pellet fraction measured by band densitometry (n=2-4 depending on the conditions). (e) Quantification of the GAC_PH_ pellet fraction measured by band densitometry (n=2-4 depending on the conditions).

### Targeted mutations in GAC PH domain are not fitness conferring *in vivo*

*T. gondii* mutant lines were generated in which we replaced the PH domain sequence at the endogenous locus was replaced with PH domain sequences containing the PA-binding mutation together with C-terminal Ty tag. Parasite lines were obtained with a wild-type PH domain (GACWT -Ty), with the individual targeted mutations (GAC^RK^ - Ty and GAC^KER^ -Ty), the double mutation patch (GAC^RKKER^ -Ty), and a line in which the PH domain was replaced by the *P. falciparum* version (GAC^PfPH^ -Ty). All GAC versions, wild-type, mutated and chimeric, localised correctly at the PCRs of the parasite as well as cytosolically (Figure 6a), suggesting that the PA-binding by the GAC PH domain is not critical for its apical localization. Likewise, upon triggering parasite motility with BIPPO, all the strains were able to display the characteristic basal accumulation of GAC (Figure 6b). Finally, to access if there was any fitness cost induced by the mutations, wild-type and mutated parasites were analysed by plaque assay and revealed that the mutations were not detrimental for the parasite lytic cycle (Figure 6c). Accordingly, no clear defect in invasion could be observed in parasites expressing the mutated versions (Figure 6d). A small reduction of the plaque size and reduction in invasion capacity was observed for the TgGAC-PfPH chimera.

**Figure 6.**
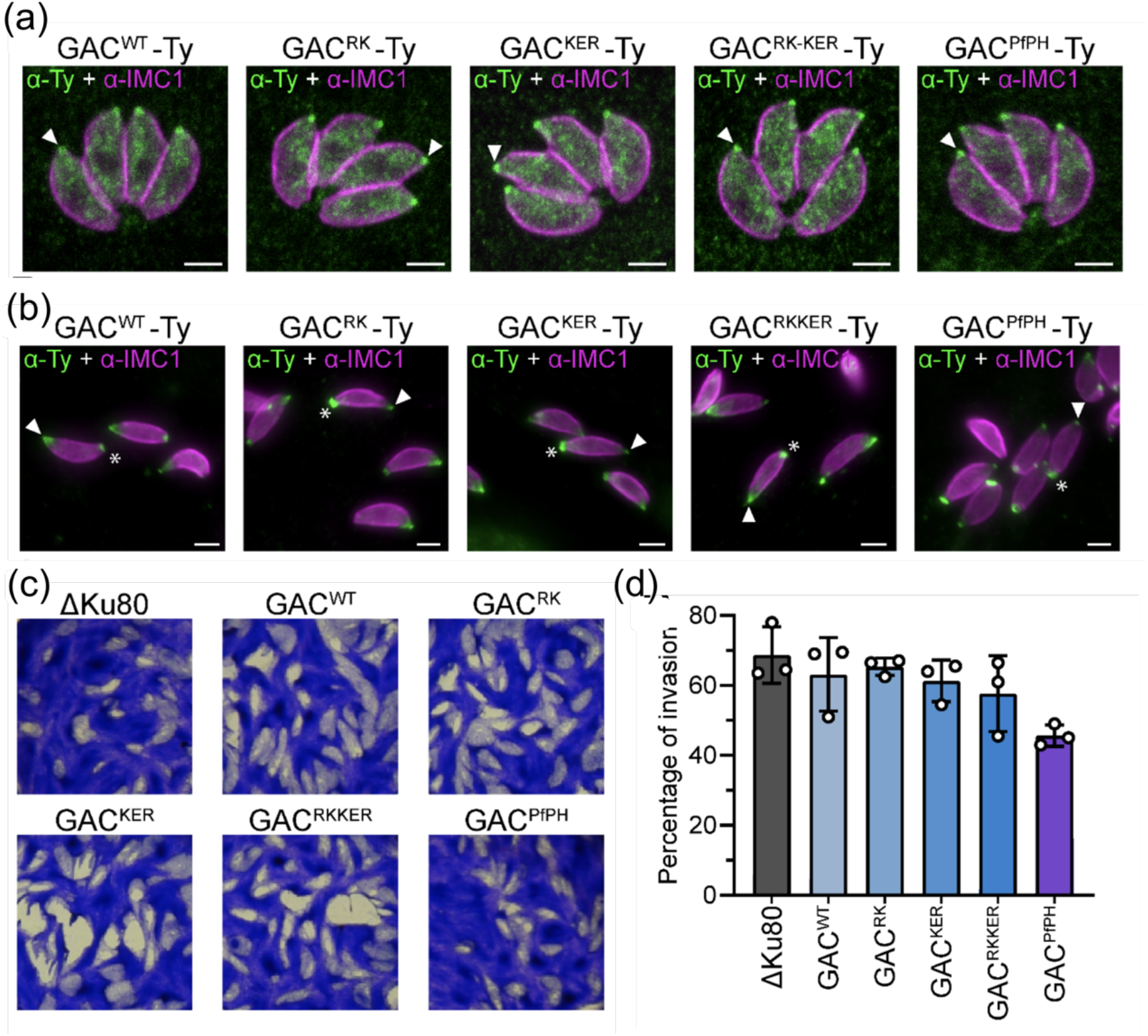
Phenotypical analysis of parasite bearing PA-binding mutations of GAC. A- GAC localization by IFA in intracellular parasites. White arrow = apical pole. Scale bar = 2μm. B- GAC localization by IFA in extracellular parasites. White arrow = apical pole. White star = basal pole. Scale bar = 2μm. C- Plaque assay of the different mutant analysed. D- Red/green invasion assay (n=3).

### Evaluation of GAC and GAC fragments binding to *toxoplasma* F-actin

Association to rabbit F-actin was previously shown to involve the N-terminal 1114 amino acids of TgGAC, roughly corresponding to two-thirds of the RI region. To examine further the interaction of TgGAC with the shorter and highly dynamic actin filaments characteristic of *T. gondii* (Skillman *et al*, 2013), a series of recombinant TgGAC fragments were purified and tested for binding. Both full-length TgGAC and a fragment encompassing residues GAC1-1114 could co-sediment with *toxoplasma* F-actin produced from TgACT (Figure 7a). In contrast, a shorter N-terminal fragment, (GAC1-617) failed to interact with F-actin. This suggests that either the binding site for F-actin lies between residues 619 to 1114 of TgGAC or that that isolated N-terminal fragment TgGAC_1-619_ cannot adopt the 3D conformation necessary for association with F-actin.

**Figure 7.**
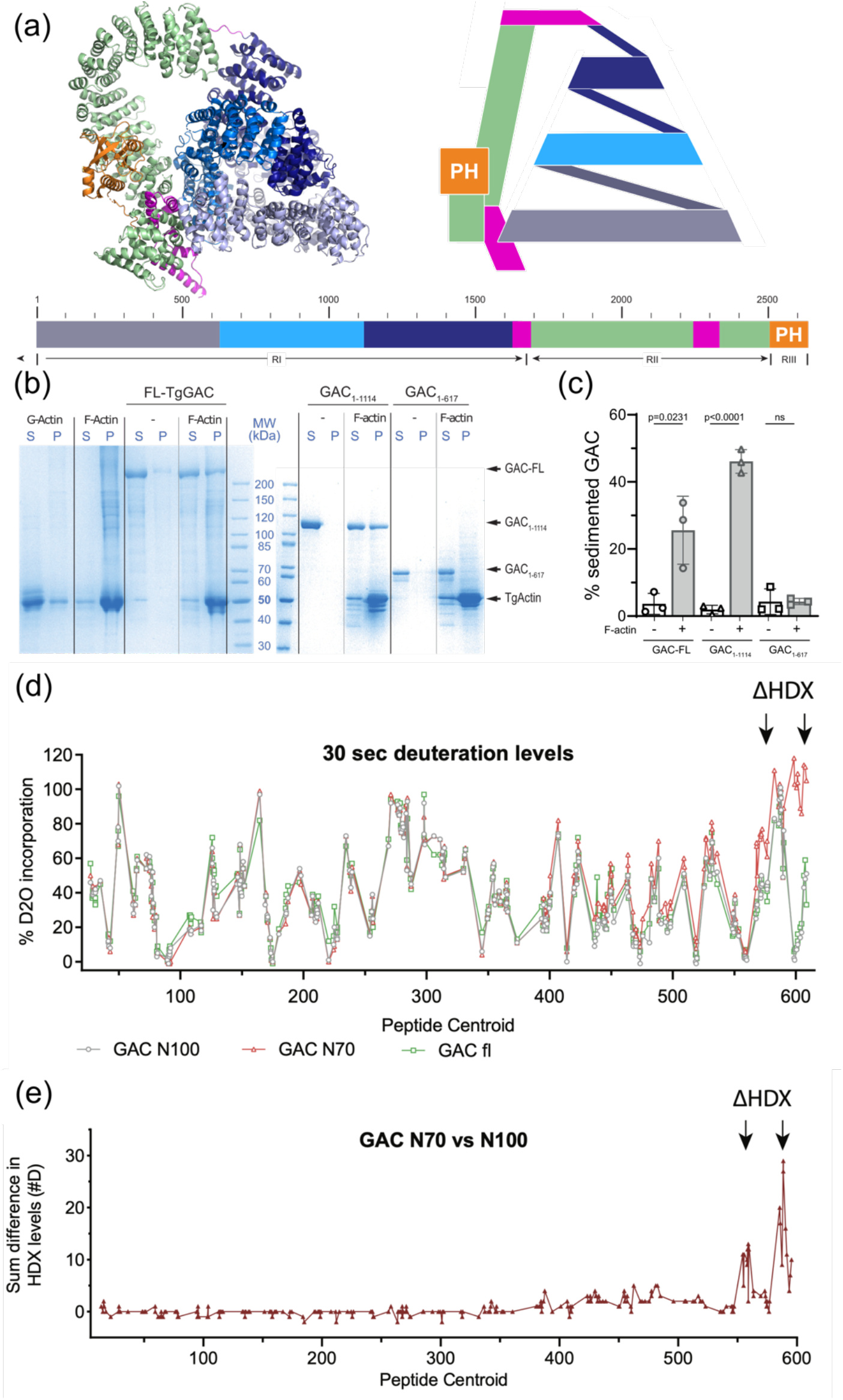
GAC interaction with TgActin filaments. (a) Schematic representation of the structure of GAC with domains colour coded (b) Coomassie-stained gel analysing proteins remaining in supernatant or sedimenting upon centrifugation at 100’000 g. S: Supernatant. P: Pellet. (c) Graphical representation of the percentage of GAC co-sedimenting alone or in the presence of TgActin filaments. Data are mean + SD of three independent experiments. P value calculated using unpaired t-test. (d) Graph comparing deuteration levels for GAC1-619, GAC1-1114 and GAC-FL. Each dot represents a single peptide where the deuteration level in percentage maximal deuteration is plotted according to the residue number at the center of the peptide. Difference plot comparing the difference in H/D exchange rate between GAC1-619 and GAC1-1114. Each dot represents a peptide with differences shown in number of incorporated deuterons.

### Insights into TgGAC conformational dynamics using hydrogen/deuterium exchange coupled to mass spectrometry

To evaluate the 3D conformations of the recombinant GAC fragments, protein dynamics was studied using hydrogen/deuterium exchange coupled to mass spectrometry (HDX-MS). HDX-MS is a powerful method that looks at protein dynamics by monitoring the exchange rate of protein amide hydrogens with the solvent (James *et al*, 2022). Analysis of peptides common to all GAC constructs shows that the vast majority of GAC N-terminus has unchanged H/D exchange rates, with only the C-terminal extremity of GAC_1-619_, encompassing residues 559-576 and 590-609, that differ from GAC 1-1114 and full-length GAC (FL-GAC) (Figures 7d, 7e, S5 and Table 2). This indicates that all fragments share a similar structure at the bottom of the pyramid (residues 1-539) and adopt a conformation mimicking the full-length enzyme. As expected, deletion of the amino acid residues following 619 leads to increased conformational flexibility within the 539-619 region, which is indicative of partial unfolding of the final AHR.

**Table 2.**
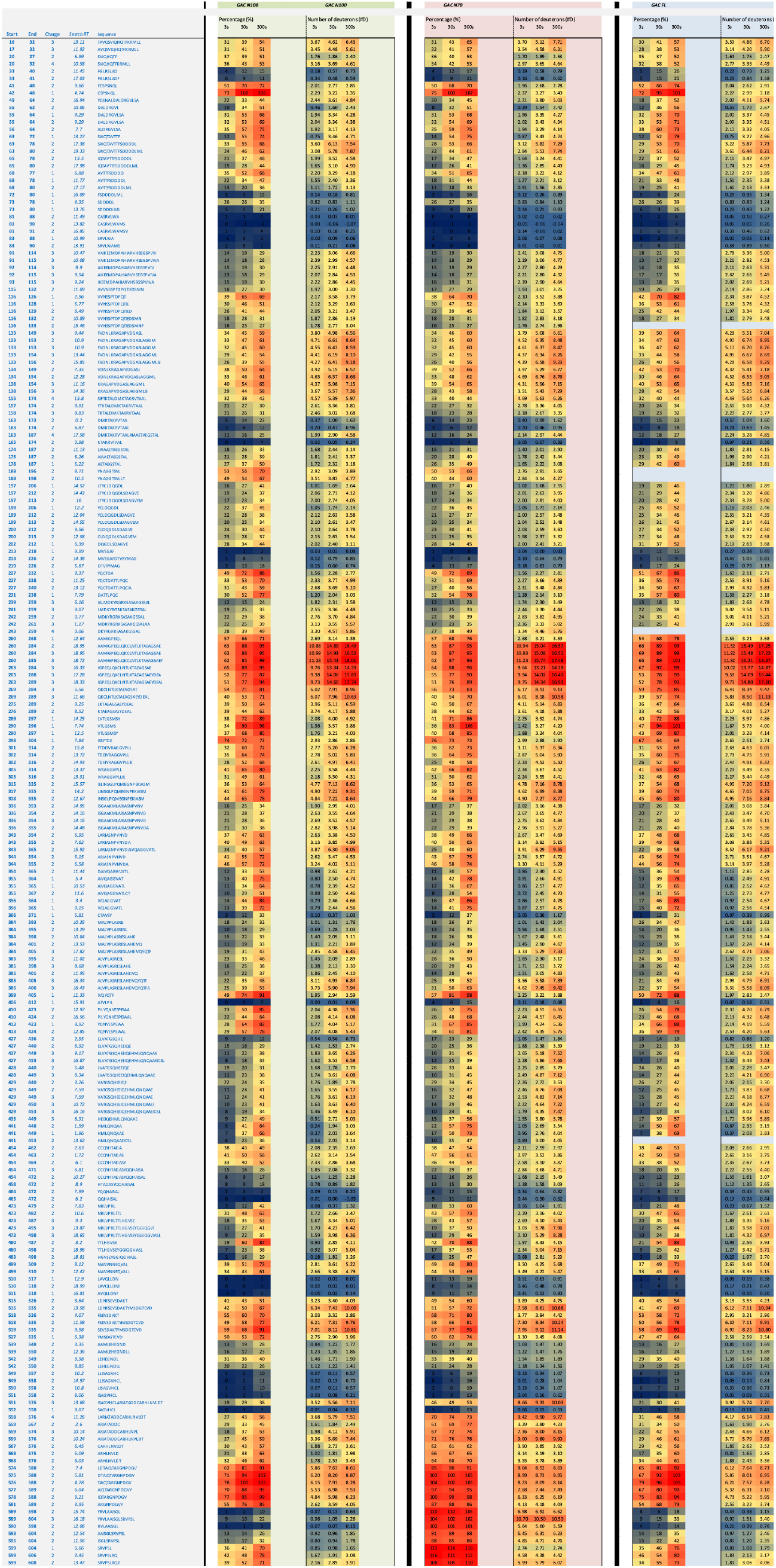
Deuterium incorporation for each of the selected peptides used for HDX-MS analysis are presented. Results are shown both as percentage deuteration compared to a theoretical maximal deuteration level and as number of deuterons incorporated into the peptide (#D)

Comparison of GAC1-1114 with FL-GAC also shows highly similar dynamics. Interestingly, FL-GAC has several peptides in the 91-133 region displaying bimodal distribution, referred to as EX1 kinetics (Weis *et al*, 2006). Bimodal distribution suggests the presence of conformational changes between two distinct populations. The N-terminal layer of the pyramid directly the C-terminal RII, so the presence of two populations within the 91-133 region can be explained by the presence of the compacted closed conformation and an open conformation, reminiscent of multiple GAC conformations in solution.

### Modelling TgGAC binding to MIC2

Intriguingly, the TgGAC PH domain contacts the C-terminal AHR arch region (AHRs 38-53), which display significant structural similarity with the armadillo repeat region (ARM) of β-catenins, superimposing with an RMSD of 4.2 Å over the backbone of 348 equivalent residues (Figure 8a). Even though the PH domain represents a minor portion of the full GAC structure, its interaction with RII region seems crucial for GAC structural integrity, as a construct lacking the PH domain cannot fold properly. The β-catenins play a strikingly similar role to GAC within adherens junctions by regulating the mechanical coupling between cells through the actin cytoskeleton and specific cellsurface adhesins (Valenta *et al*, 2012). β-catenins bind extended cadherin tails emerging from the plasma membrane via a superhelical surface formed by 12 tandem ARM repeats (Choi *et al*, 2009; Huber & Weis, 2001; Ishiyama *et al*, 2010). To explore this, AlphaFold2, which has recently been shown to perform well for the prediction of protein-peptide complexes (Choi *et al*., 2009; Jumper *et al*., 2021), was used to generate a model for the complex of TgGAC1670-2639 with the C-terminal 20 amino acids residues of MIC2. In this model, the MIC2 tail binds the concaved surface formed by AHR39-46 (Figure 8a), which coincides well with interface formed in β-catenin/E-cadherin complexes suggesting a role for this ARM arch structure in recruiting MIC2 to the glideosome machinery.

**Figure 8.**
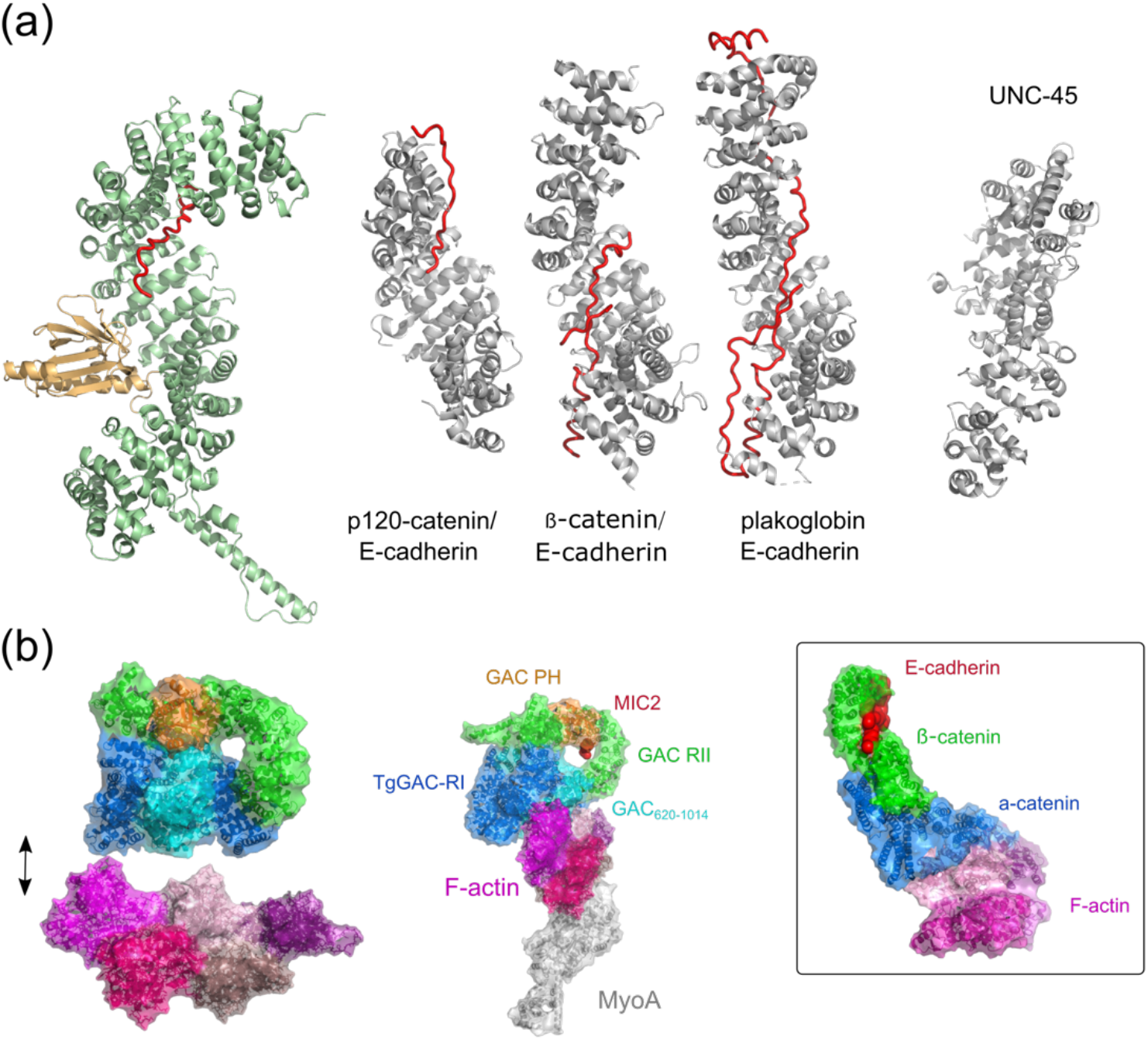
Structural homology for TgGAC and glideosome model. (a). Superposition of model of the TgGAC/MIC tail complex with human p120 catenin/E-cadherin (PDB 3L6X), β-catenin//E-cadherin from *Caenorhabditis elegans* (PDB 4R10), human plakoglobin//E-cadherin (PDB 3IFQ) and UNC45 myosin chaperone *Caenorhabditis elegans* (PDB 6QDL), (b) Model for GAC within the glideosome

## Discussion

Apicomplexan parasites, for which *T. gondii* is a model organism, propel themself by a specialized actomyosin-dependent gliding motility that relies on a large, conserved protein to connect actin filaments with parasite plasma membrane. GAC, an abundant protein that crucially localizes to the PCRs at the apical tip of parasite was shown to adopt multiple conformations to regulate its function. The most striking feature of the TgGAC structure is the large continuous supercoiled ARM region (residues 1-2490) which forms a ring. The closure of the ring results from an extensive interface between the ARM1 and ARMs14-16 from RI and ARMs46-48 from RII (N/C-interface - Figure 2). This closed conformation was unexpected as an earlier SAXS model suggested an extended, club shape molecule in solution with no evidence for a N/C terminal interface (Jacot *et al*., 2016). Despite this discrepancy, the interface formed in the closed structure is highly conserved at the protein sequence level suggesting that it is a functionally relevant state. Subsequent pH dependent SAXS analyses reported in the present study revealed that an open structure exists at high pH, but oligomerised forms predominate at low pH. The nature of the N/C interface in the closed structure is hydrophilic with several electrostatic interactions, including some that are readily titratable, such H24 and H2164. It is conceivable that deprotonation of these residues at high pH removes key salt bridges that stabilise the closed conformation and facilitate a conformationally labile open structure. Our SAXS data at low pH provide some support for this notion, as it reveals a multimerised state. While this is unlikely to reflect oligomerisation *in vivo*, it could result from the reformation of these electrostatic interactions in an intermolecular manner. The ability to adopt both a flexible open conformation and transition to a compact closed structure could play role in assisting GACs assembly on the glideosome and at the parasite membrane.

GAC interacts with the PA-enriched membranes that are generated during motility signalling pathways. The C-terminal PH domain within GAC presents a contiguous, positively charged surface and while this is important for PA binding *in vitro*, its mutation does not cause a significant defect in GAC localisation, translocation, or fitness. Other regions in GAC, such as the basic protrusion in AHR53 (Figure 4), are also likely to contribute to membrane binding *in vivo*. Furthermore, the presence of a membrane interaction surface within the TgGAC PH domain suggests that the binding site for the juxtamembrane region of the MIC2 cytoplasmic tail is located nearby and these interactions may also be cooperative. Even though the PH domain represents a minor portion of the full-length GAC structure, its interaction with RII region seems crucial for GAC structural integrity, as a construct lacking the PH domain cannot fold properly.

The principal *toxoplasma* F-actin binding region was localised to the first 1114 residues of TgGAC (TgGAC_1-1014_), which forms a large, supercoiled base of the N-terminal pyramid (TgGAC-R1:1-1656 – Figure 2) and provides a platform with several potential sites of interaction with a helical actin filament. A shorter fragment encompassing only the first turn of the supercoiled pyramid (TgGAC_1-619_) does not interact with F-actin. Analysis of protein conformation by HDX-MS confirmed that TgGAC_1-619_ adopts a similar conformation as TgGAC, suggesting that the actin-binding interface lies between residues 620-1114 which starts the second supercoil turn. The absence of Factin binding for the N-terminal portion, which forms the base of GAC pyramidal structure, suggests a role this region in stabilizing the closed conformation by making direct contacts with the RII region. Membrane association simulations for TgGAC reveal a specific membrane binding surface involving the PH domain and RII, and this orientation places the TgGAC_1-619_ region distal for interaction with the actin filament (Figure 8b).

As the available space between the parasite inner membrane complex and the plasma membrane is insufficient for the open structure of GAC (i.e. with RII & RIII extended) to bridge F-actin to the plasma membrane lengthways, the closed structure is likely to represent a functional important state when GAC is fully engaged within the glideosome. Other actin-membrane bridging proteins, like the ERM (ezrin, moesin, and radixin) family of proteins, crosslink cortical actin to plasma membrane and full engagement is achieved by a reorganisation of actin-binding regions by cooperative interactions with phosphatidylinositol 4,5-bisphosphate (PIP2) (Ben-Aissa *et al*, 2012). It is therefore conceivable that PA binding and MIC2 recruitment at the plasma membrane by GAC also contributes to stabilising the closed conformation.

The function of GAC in bridging parasite cytoskeleton to the host cell substrate is reminiscent of that for the mammalian catenins within adherens junctions, which comprise several components (Figure 8b). Nominally, α-catenin crosslinks F-actin to β-catenin, while β-catenin establishes the connection to the E-cadherin tails. The single chain of GAC carries out both these roles, i.e. RI acts like the actin-binding α-catenin and is linked via L1 to RII ARM arch structure, which like β-catenin tethers the system to the cell surface adhesins. Intriguingly, the extensive interface within and between the catenins and E-cadherin facilitates molecular transduction via mechanical strain (Angulo-Urarte *et al*, 2020; Bush *et al*, 2019). Mechanical force induces a rearrangement of binding interfaces that results in a strengthening of its interaction with F-actin and this effect is dependent on the direction of applied force (Mei *et al*, 2020; Xu *et al*, 2020). Such catch bond behaviour may also be the relevant for GAC function. Importantly, part of the 620-1114 F-actin binding region is not fully accessible in the closed conformation, therefore is also conceivable that optimal F-actin binding requires structural rearrangement. The stabilised closed conformation would be able to both resist and sense the significant inter parasite-substrate forces generated by MyoA translocation of F-actin. Such force acting along parasite–host cell interface could open additional cryptic binding sites that strengthen F-actin binding and ensure a coordinated direction of motion.

Structural similarity also exists between the C-terminal ARM region of GAC (TgGAC_1670-2639_) and the family of myosin-specific chaperones which possess ARM-rich UCS (UNC-45/Cro1/She4) domains (Hellerschmied & Clausen, 2014) that interact with myosin motors domains (Figure 8a). While *T. gondii* possesses a dedicated UCS chaperone for TgMyoA (TgUNC) (Bookwalter *et al*, 2014), which is critical for successful folding of the motor domain (Frénal *et al*, 2017b), the similarity between the GAC C-terminal ARM region and the UCS chaperones is intriguing. Myosin chaperones also have the propensity to multimerise and in some cases form chains that assist myosin assembly on the filament (Gazda *et al*, 2013). Although no direct evidence for an interaction between TgGAC and TgMyoA has been found, it is tempting to speculate that TgGAC may assist TgMyoA organisation on F-actin.

## Materials and methods

### Protein expression and purification

Full-length TgGAC gene with TEV cleavable N-terminal 6xHis-tag has been cloned into the pET28a vector as previous described (Jacot *et al*., 2016; Kumar *et al*., 2022). Constructs for PfGAC_PH_ (PfGAC_2471-2605_) and TgGAC_PH_ (TgGAC_2505-2639_) were construct with a 6His purification tags and an additional SUMO tag for soluble expression of PfGAC_PH_. A Q5 site-directed mutagenesis kit (NEB) and the manufacturers protocol were used to generate and PfGAC_PH_ mutants with standard primers. For protein expression, plasmids were transformed into BL21 (DE3) (NEB) or Rosetta2 (Novagen) *E. coli* strains. Expression was carried in minimal medium supplemented with ^15^NH4Cl and/or ^13^C-glucose for NMR isotopic labelling. Purification for His-tagged constructs was carried by Ni^2+^ affinity chromatography. Removal of the SUMO tag for PfGAC_PH_ samples was carried out by incubation with purified ULP1 protease. Further purification for all constructs was achieved with by size exclusion chromatography.

### NMR spectroscopy

Samples of purified ^15^N/^13^C-PfGAC_PH_ and ^15^N/^13^C-TgGAC_PH_ were prepared and supplemented with D20. All NMR spectra were acquired at 298K on Bruker Avance-III DRX 800 and Avance-III 600 spectrometers. Triple resonance HNCA, HNCACB, HNCO and HN(CO)CA spectra (Sattler *et al*, 1999) were recorded and analysed to obtain backbone assignments, which was assisted using MARS program (Jung & Zweckstetter, 2004). Chemical shift assignment and analysis was performed using an in-house version of NMRview (Marchant *et al*, 2008). Dihedral angles were calculated using TALOS+ (Shen *et al*, 2009).

For PfGAC_PH_ and TgGAC_PH_ 1D ^1^H NMR LUV binding assays performed with LUVs containing an increasing proportion of POPA (POPA Mol% value). For each LUV composition titration series, a separate Kdapp value was calculated for each replicate (n=3), and a mean Kdapp value calculated. PfGAC_PH_ and TgGAC_PH_ titration PRE experiments were carried out with increasingly POPA-enriched MSP1D14_-_5 nanodiscs Relative signal reductions for selected residues were determined and napped.

### X-ray data collection and processing

Diffraction data from a single native crystal were collected on beamline i04 of the Diamond Light Source (DLS), UK. Data were processed with CCP4, dials (Beilsten-Edmands *et al*, 2020; Winn *et al*, 2011; Winter, 2010; Winter *et al*, 2018) and scaled using dials.scale (Evans, 2006) within the Xia2 package (Winter *et al*, 2013). Multiple-wavelength anomalous diffraction (MAD) data from a single SeMet labelled crystal were collected on beamline i04 of the Diamond Light Source at the following wavelengths: peak=0.9795Å, inflection=0.9796Å and remote=0.9722Å. Data were processed initially by AutoProc (Vonrhein *et al*, 2011). Substructure definition and initial model building were performed using AutoSHARP (Vonrhein *et al*, 2007). This was followed by manual building in Coot (Emsley *et al*, 2010) and further refinement using Phenix Refine (Adams *et al*, 2010). Data collection statistics have been published previously (Kumar *et al*., 2022). The structure has been deposited in wwPDB under accession code: PDB ID 8C4A (Berman *et al*, 2007).

### Coarse grained (cgMD) simulations

The GAC crystal structure was rotated randomly in respect to the membrane patch to generate the first set of coordinates. For two independent replicates GAC was rotated again (0, 90) and (90, 0). CHARMM-GUI was used to generate a system in which GAC was translated 12 nm in the z axis from the centre of the membrane. The membrane composition used was 50% POPC: 50% POPA, to replicate simulations performed in Darvill *et. al*. (Darvill *et al*., 2018). Simulations were performed using the gromacs biomolecular software package and the MARTINI 3 forcefield with ELNEDYN restraints. Simulations were performed at 303.15 K, using the V-rescale algorithm with a tau 1.0. Production simulations were 10 μs in length. Analysis was performed using gromacs tools, VMD, and the ProLint server.

### Small angle X-ray scattering (SAXS)

Samples for SAXS measurements were prepared by concentrating samples from SEC using a centrifugal spin device with a molecular weight cutoff of 100 kDa. For all experiments a buffer of 25 mM Tris, 5 mM TCEP, pH 8.0 was used. SAXS data were measured using a laboratory based flux-optimized Bruker AXS Nanostar with a gallium liquid metal jet source (Schwamberger *et al*, 2015) and scatterless slits (Li *et al*, 2008). More information about the optimized instrument can be found here (Lyngso & Pedersen, 2021). All data were measured for 1800 s at 20 °C. SAXS data are plotted as intensity as a function of *q*, which is the modulus of the scattering vector and if defined as *q* = (4πsin(θ))/λ_Ga_, where 2Θ is the scattering angle between the incident and scattered beam and λ_Ga_ = 1.34 Å. Data were background subtracted and converted to absolute scale using the software package SUPERSAXS (CLP Oliveira and JS Pedersen, unpublished). The mass was calculated using 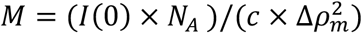, where *I*(0) is the intensity extrapolated to *q* = 0, *N*_A_ is Avogadro’s number, *c* is the protein concentration is mg/mL, and Δ*p_m_* is the scattering contrast per mass that can be estimated to 2.0 × 10^10^ cm/g for a typical protein. *I(0)* was determined both by a Guinier fit analysis (using the intercept with the y-axis) and from an indirect Fourier transform (IFT) routine (Glatter, 1977). The theoretical scattering curve was calculated using the program wlsq_PDBx (Steiner *et al*, 2018). An *ab initio* reconstruction of the protein structure was performed using GASBOR from the ATSAS package (Svergun *et al*., 2001) where the number of amino acids is given, and each amino acid is represented by a dummy residue. The optimization was performed with the real space option due to the large size of GAC.

### Molecular dynamics simulations

The TgGAC crystal structure was used to generate coarse-grained protein: membrane simulation systems in which TgGAC was rotated randomly in respect to the membrane patch to generate the first set of coordinates. CHARMM-GUI Martini Maker was used to generate a system in which GAC was translated 12 nm in the z axis from the centre of the membrane (Qi *et al*, 2015). The membrane composition used was 50% POPC: 50% POPA, to replicate simulations performed in Darvill et al. (Darvill *et al*., 2018). For two independent replicates TgGAC was rotated again (0, 90) and (90, 0). Simulations were performed using the gromacs biomolecular software package version 2021.3 (Hess *et al*, 2008)and the MARTINI3 forcefield with ElNeDyn restraints (Souza *et al*, 2021). The v-rescale thermostat (tau 1.0 ps) (Bussi *et al*, 2007) and the Parrinello–Rahman barostat (tau 12.0 ps) (Parrinello & Rahman, 1981)were used to maintain temperature (303.15 K) and pressure (1 bar). Production simulations were 10 μs in length. Analysis was performed using gromacs tools, VMD, (Humphrey *et al*, 1996) and the ProLint server (Sejdiu & Tieleman, 2021).

### Plaque assay

HFFs monolayers were infected with freshly egressed parasites and incubated for 7 days at 37°C. Cells were then fixed with 4% PFA / 0,05% Glutaraldehyde for 10-15 minutes. After neutralization with 0,1M glycine/PBS, cells were stained using crystal violet. For quantification, pictures were taken, and the plaque area was determined using ImageJ.

### Immunofluorescence assay

For vacuoles images, parasites were inoculated on an HFF monolayers previously seeded on a glass coverslip. The parasites were grown for 16-24h at 37°C. For extracellular parasites, freshly egressed parasites were seeded on gelatin-coated coverslips and media containing BIIPO was used to stimulate motility. The coverslips were then fixed with 4% Paraformaldehyde (PFA) / 0,05% Glutaraldehyde (PFA-Glu) during 10 min. The fixative agent was then neutralized using 0,1M glycine/PBS for 10 min. Cells were then permeabilized for 20 minutes using 0,2% TritonX100/PBS and blocked using 5% BSA/PBS for 20 minutes before incubation with primary antibodies in 2% BSA/0,2% TritonX100/PBS for 1 hour. Coverslips were then washed three times for 5 minutes using 0,2% TritonX100/PBS. Secondary antibodies were incubated 1 hour similarly to the primary antibodies. Finally, the coverslips were washed three times with PBS before mounting on glass slides using DAPI-containing Fluoromount. For immunofluorescence analysis, the secondary antibodies Alexa Fluor 488 and Alexa Fluor 594 conjugated goat α-mouse/rabbit antibodies (Molecular Probes) were used.

### Liposome binding assay

Proteins were expressed in bacteria and purified by the PPR2P platform (University of Geneva). The liposomes were prepared in-house as follow: lipids were mixed in glass vials with the following proportions: 10% DOPE + X%DO-PA + qsp%DOPC (percentage by weight). The mix were slowly dried using nitrogen flow. The lipids were dried further in a dessicator for 30min. The dried lipids were then resuspended in “lipid buffer” (50mM Hepes pH 7.5 / 100mM NaCl / 5% Glycerol) to reach a 5 mg/mL concentration. Resuspension was then ensured by vortexing the mixes for 5min. Then, seven freeze-thaw cycles were performed (20sec in liquid nitrogen followed by 90sec in a 33°C water-bath). Lipid mixes were extruded by passing them through 0,1μm filters 21 times. Liposomes were then aliquoted in microtubes (50μl), flash frozen in liquid nitrogen, and stored at −80°C until use.

For the binding assay, proteins were resuspended in “protein buffer” (20 mM Tris pH 7.4 / 200 mM NaCl / 1 mM DTT), centrifuged at 100,000g for 30 min at 4°C to remove any precipitate and measure the exact protein concentration in the supernatant. Protein concentration was then adjusted to 0.2 mg/mL using protein buffer. In parallel, the liposomes were diluted at 2 mg/mL in “lipid buffer” (50 mM Hepes pH 7.4/ 100 mM NaCl/ 1% Glycerol) and re-diluted two-fold using water to achieve a concentration of 1mg/ml. For the “no lipid” conditions, lipid buffer was simply diluted two-fold in water. For the reaction, 40 μL of protein and 40 μL of liposomes were mixed and incubated on ice for 1 hour. The mix was then centrifuged at 120,000g for 30 minutes at 4°C. Supernatant and pellets were separated, resuspended in Laemmli buffer, and boiled 10 min at 95°C. Equal volumes were then ran on polyacrylamide gels and stained using Coomassie blue. Quantification was performed by band densitometry.

### Strains generation – GAC mutations at the endogenous locus

The pGST-GAC_PH domain vector used to produce the PH domain of GAC was used as a base to generate the points mutation. The points mutations were inserted using the Q5 mutagenesis kit (New England Biolabs) and specific primers. The mutations in the PH domains were checked by sequencing. Then, the 5’UTR_UPRT-pT8-DDGACΔPH-Ty-DHFR-3’UTR_UPRT was linearized with SgrAI while the mutated PH domain from the pGST vector was amplified by PCR using primers containing homology regions. The mutated PH domain was inserted in the linearized 5’UTR_UPRT-pT8-DDGACΔPH-Ty-DHFR-3’UTR_UPRT by Gibson assembly. The insertion of the PH domain in the receiving vector was checked by analytical digestion and sequencing. The mutated PH domain of newly generated 5’UTR_UPRT-pT8-DDGACmut-Ty-DHFR-3’UTR_UPRT vector (therefore without introns), was amplified by KOD PCR using specific primers containing homology regions (on one side with the sequence preceding the PH domain, and on the other with the 3’UTR of the GAC gene). A double gRNA targeting the region preceding the PH domain and the 3’UTR of the GAC gene was generated in parallel by Q5 mutagenesis. For transfection, the equivalent of 100μl of KOD PCR and 40μg of gRNA was used. 48h after transfection, the parasites were FACS sorted to select clones expressing the gRNA-Cas9-YFP construct. Then, the individual clones were amplified, genomic DNA was extracted, and integration PCR of the GAC PH domain region was checked (the PCR ensured the loss of the introns, generating a smaller PCR amplicon in modified parasites compared to wild types). In addition, the PH domain of the PCR positive clones were fully sequenced.

### HDX-MS sample preparation and data analysis

HDX-MS experiments were performed at the UniGe Protein Platform (University of Geneva, Switzerland) following a well-established protocol with minimal modifications (Wang *et al*, 2018). Details of reaction conditions and all data are presented in Tables S1 and S2. HDX reactions were done in 50 μl volumes with a final protein concentration of 2.4 μM of GAC protein. Briefly, 120 picomoles of protein in 10 μl final volume were pre-incubated 5 min at 22 °C before the reaction.

Deuterium exchange reaction was initiated by adding 40 μl of D2O exchange buffer (20 mM Tris pH 8, 15 0mM NaCl, 5 mM DTT in D2O) to the protein-peptide mixture. Reactions were carried-out at room temperature for three incubation times (3 s, 30 s and 300 s) and terminated by the sequential addition of 20 μl of ice-cold quench buffer 1 (4 M Gdn-HCl, 1 M NaCl, 0.1 M Nah_2_PO4 pH 2.5, 1 % formic acid (FA), 200 mM TCEP). Samples were immediately frozen in liquid nitrogen and stored at −80 °C for up to two weeks. All experiments were repeated in triplicate.

To quantify deuterium uptake into the protein, samples were thawed and injected in a UPLC system immersed in ice with 0.1 % FA as liquid phase. The protein was digested via two immobilized pepsin columns (Thermo #23131), and peptides were collected onto a VanGuard precolumn trap (Waters). The trap was subsequently eluted, and peptides separated with a C18, 300Å, 1.7 μm particle size Fortis Bio 100 x 2.1 mm column over a gradient of 8 – 30 % buffer C over 20 min at 150 μl/min (Buffer B: 0.1% formic acid; buffer C: 100% acetonitrile). Mass spectra were acquired on an Orbitrap Velos Pro (Thermo), for ions from 400 to 2200 m/z using an electrospray ionization source operated at 300 °C, 5 kV of ion spray voltage. Peptides were identified by data-dependent acquisition of a non-deuterated sample after MS/ MS and data were analyzed by Mascot. All peptides analysed are shown is Tables S1 and S2. Deuterium incorporation levels were quantified using HD examiner software (Sierra Analytics), and quality of every peptide was checked manually. Results are presented as percentage of maximal deuteration compared to theoretical maximal deuteration. Changes in deuteration level between two states were considered significant if >20% and >2 Da and p< 0.01 (unpaired t-test).

## Materials and Data availability

Reagents generated in this study will be made available upon request. All data generated or analysed during this study are included in the manuscript and supporting files. The structure has been deposited in wwPDB under accession code: PDB ID 8C4A (Berman *et al*., 2007). The mass spectrometry proteomics data have been deposited to the ProteomeXchange Consortium via the PRIDE (Perez-Riverol *et al*, 2022) partner repository with the dataset identifier PXD039335.

## Acknowledgements

This work was supported by a BBSRC and Leverhulme Trust awards to SJM (RPG_2018_107), the Independent Research Fund Denmark through grants 8021-00133B (J.S.P.) and the Swiss National Science Foundation to DSF (10030_185325 and CRSII5_198545). The crystallisation facility at Imperial College London is supported by the Biotechnology and Biological Sciences Research Council (BB/D524840/1) and Wellcome Trust (202926/Z/16/Z). We are grateful to staff at Diamond Light Source beamline I04 for their help with data collection We are thankful to Rémy Visentin at the Protein Platform of the University of Geneva for assistance with recombinant protein purification.

## Source Data

**Maps and Coordinates** wwPDB X-ray Structure Validation; MTZ file for TgGAC X-ray diffraction data and Final refined PDB file for TgGAC from X-ray diffraction

**Figure 1-source data 1:** PDB file for validated model of full-length TgGAC in the closed conformation

**Figure 2-source data 1:** Raw SAXS data for TgGAC

**Figure 3-source data 1:** Molecular dynamics trajectory analysis data for TgGAC

**Figure 4-source data 1:** Raw and processed NMR data for 1D SUV titration with PfGACPH

**Figure 4-source data 2:** Raw and processed NMR data for PRE measurements of PfGACPH with nanodiscs

**Figure 5-source data 1:** Full, uncropped SDS-page images for liposome binding experiments

**Figure 5-source data 2:** Figure and caption for Liposome binding assays using uncropped SDS-page images with key bands labelled

**Figure 5-source data 3:** Liposome binding data tables

**Figure 6-source data 1:** Parasite invasion data tables

**Figure 7-source data 1:** Full, uncropped SDS-page images for F-actin binding experiments

**Figure 5-source data 2:** Figure and caption for Actin binding assays using uncropped SDS-page images with key bands labelled

**Figure S1-source data 1:** Table of NMR assignments for TgGAC_PH_

**Figure S4-source data 1:** Table of NMR assignments for PfGAC_PH_

**Figure S4-source data 2:** Raw and processed NMR data for PRE measurements of TgGACPH with nanodiscs

---

**Figure S1.**
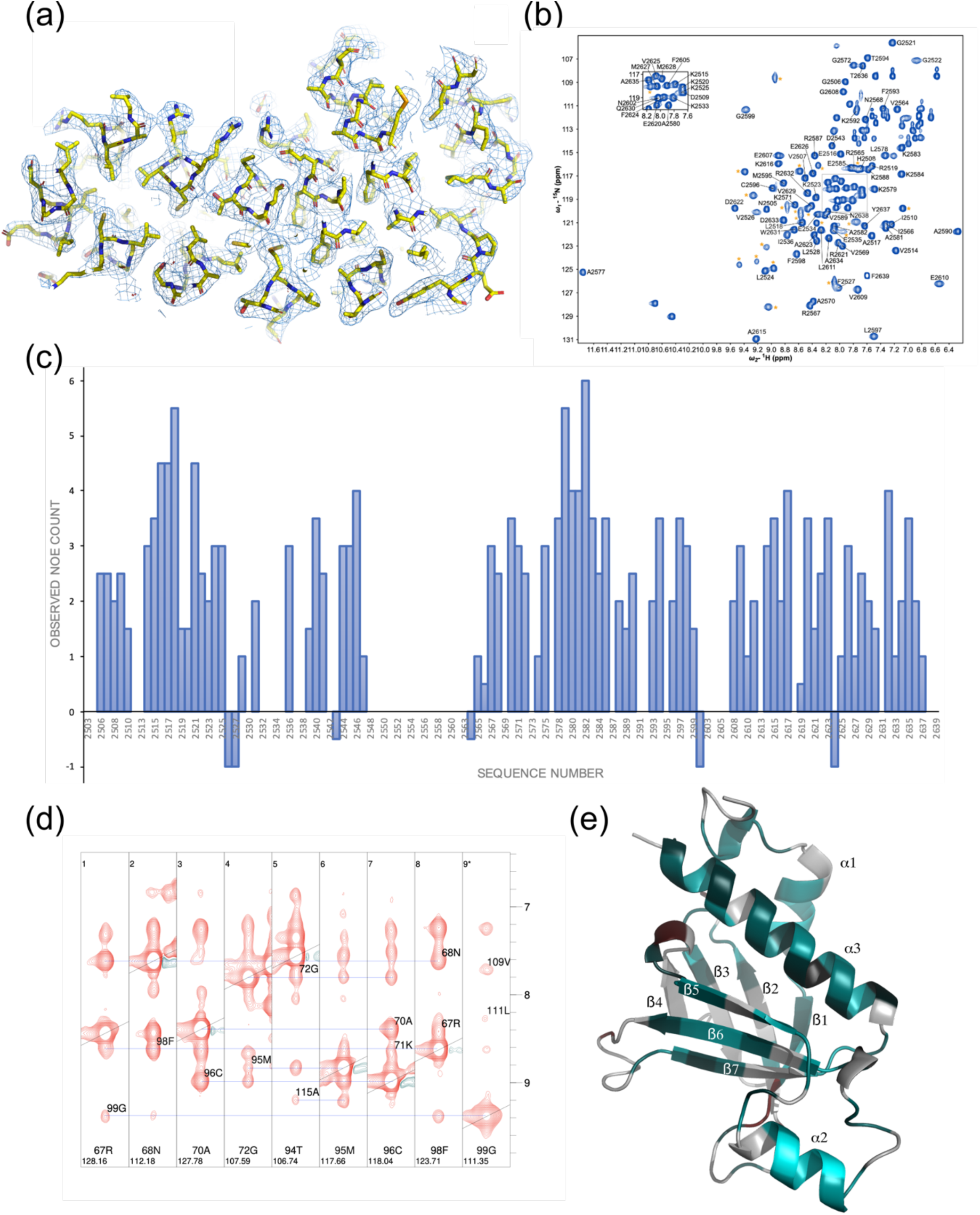
Assessing the quality of structural data for GAC. (a) Representative X-ray crystallographic electron density map showing the fit to residues 1203-1453 from the final model. (b) Assigned ^1^H-^15^N HSQC NMR spectrum for TgGAC_PH_ (TgGAC_2505-2639_), (c) Bar chart of observed inter-residue NOE count based on the Alphafold2 structure with a 5Å cutoff against sequence numbering for TgGAC_PH_. Gaps represent unassigned regions (d) Selection from strip plot showing example inter β-strand NOEs observed that are expected with the Alphafold2 structure of TgGAC_PH_. (e) Structural model for TgGAC_PH_ coloured according to the observed NOE count based on the Alphafold2 structure with a 5Å cutoff.

**Figure S2.**
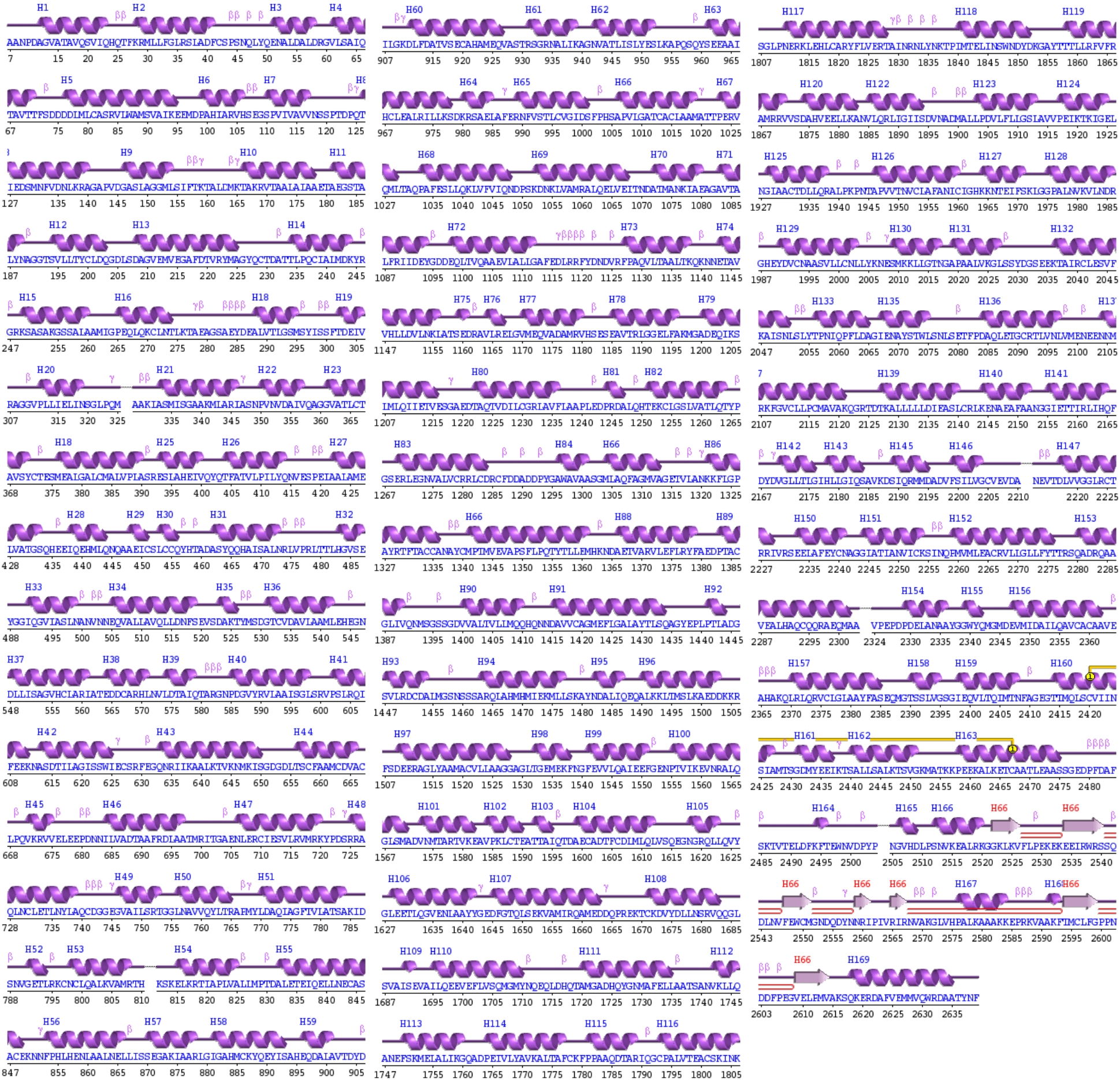
Secondary structure for TgGAC. Wired representation generated using PDBsum (Laskowski, PDBsum1: A standalone program for generating PDBsum analyses. *Protein Sci* 31, e4473 (2022). Helices labelled H1 and strands by their sheets A, B. Motifs for β-turn, g-turn and β-hairpins are labelled.

**Figure S3.**
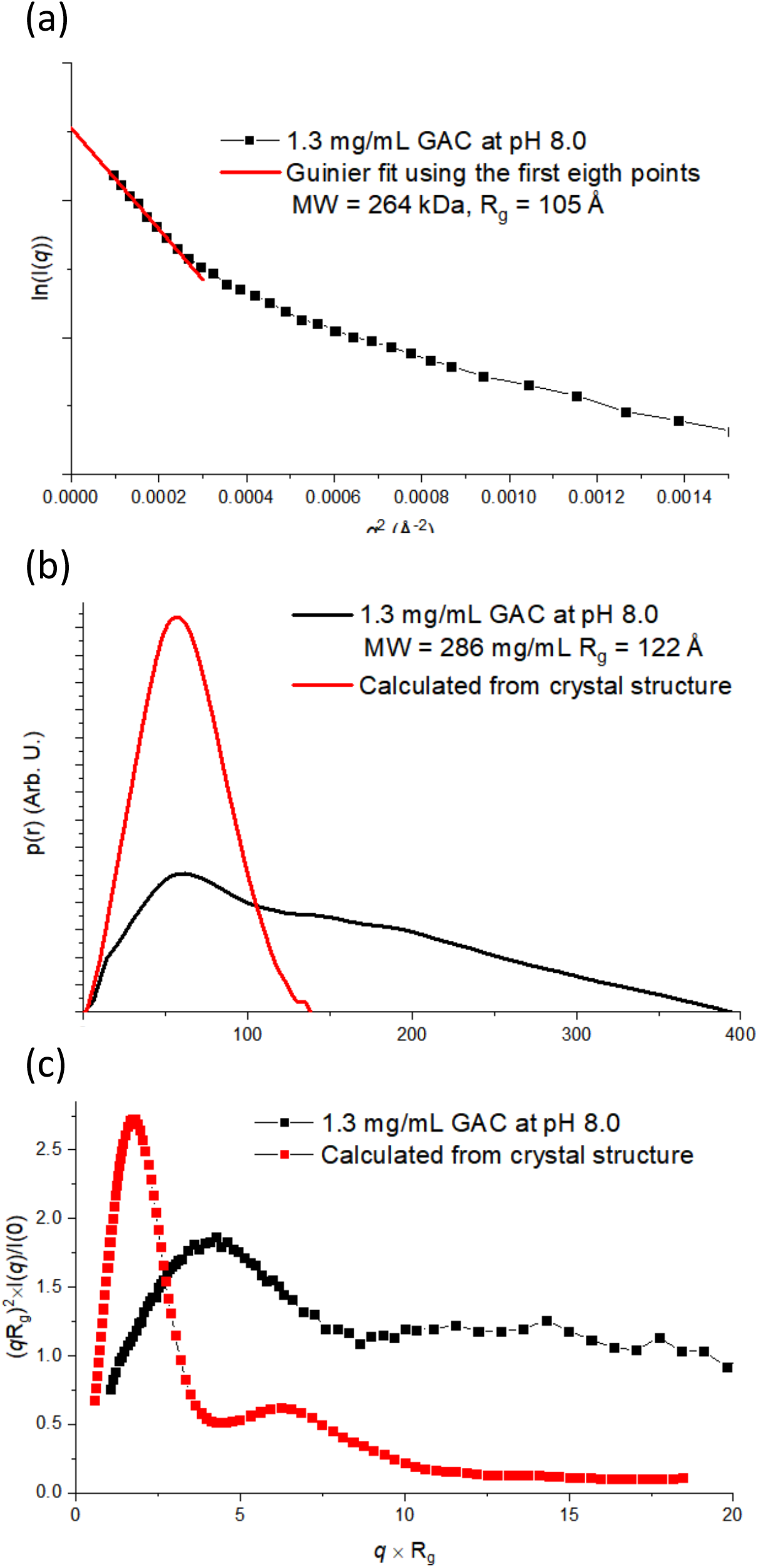
Model independent analysis of GAC SAXS data at pH 8.0 and comparison to crystal structure. (a) Guinier fit of GAC SAXS data using the first eight points. The molecular mass (MW) and radius of gyration (Rg) calculated from the Guinier fit are noted in the legend. (b) p(r) function for GAC SAXS data and calculated from crystal structure. The MW and Rg calculated from IFT routine for SAXS data are noted in the legend. (c) Dimensionless Kratky plot for SAXS data and calculated from crystal structure.

**Figure S4.**
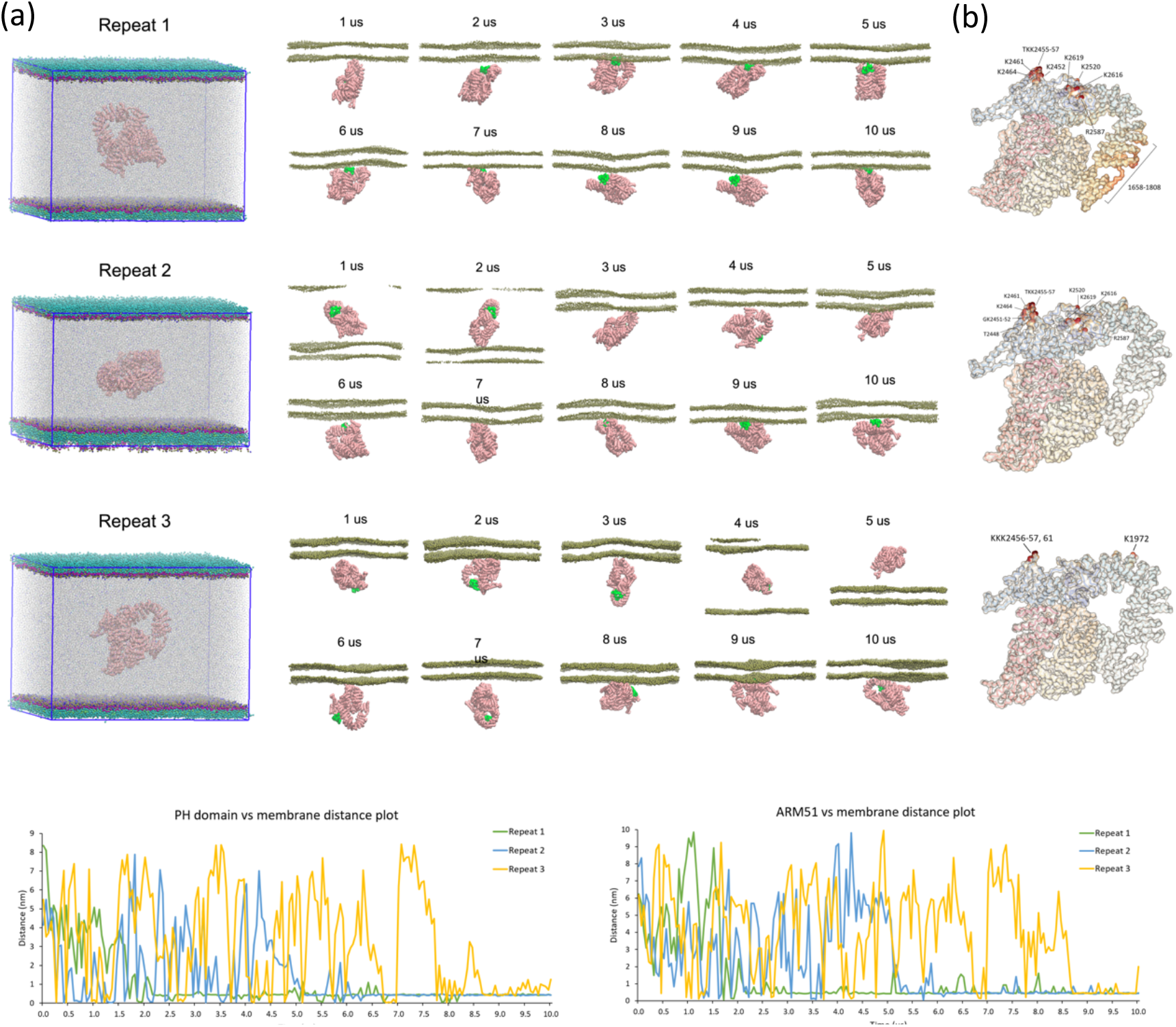
Molecular simulations of GAC and PA-containing membranes. (a) Binding series and initial orientations for each of 3 independently generated repeats. The initial configurations show GAC randomly orientated relative to the membrane patch and the corresponding time series with snapshots at each microsecond are shown. Water is omitted for clarity. (b) Lipid contact analysis for the final microsecond of each trajectory. The heatmap and labelled residues show key regions in contact with POPA lipid. (c) Centre of mass distance over time for each simulation repeat for the PH domain (left hand panel) and the ARM51 region (right hand panel).

**Figure S4.**
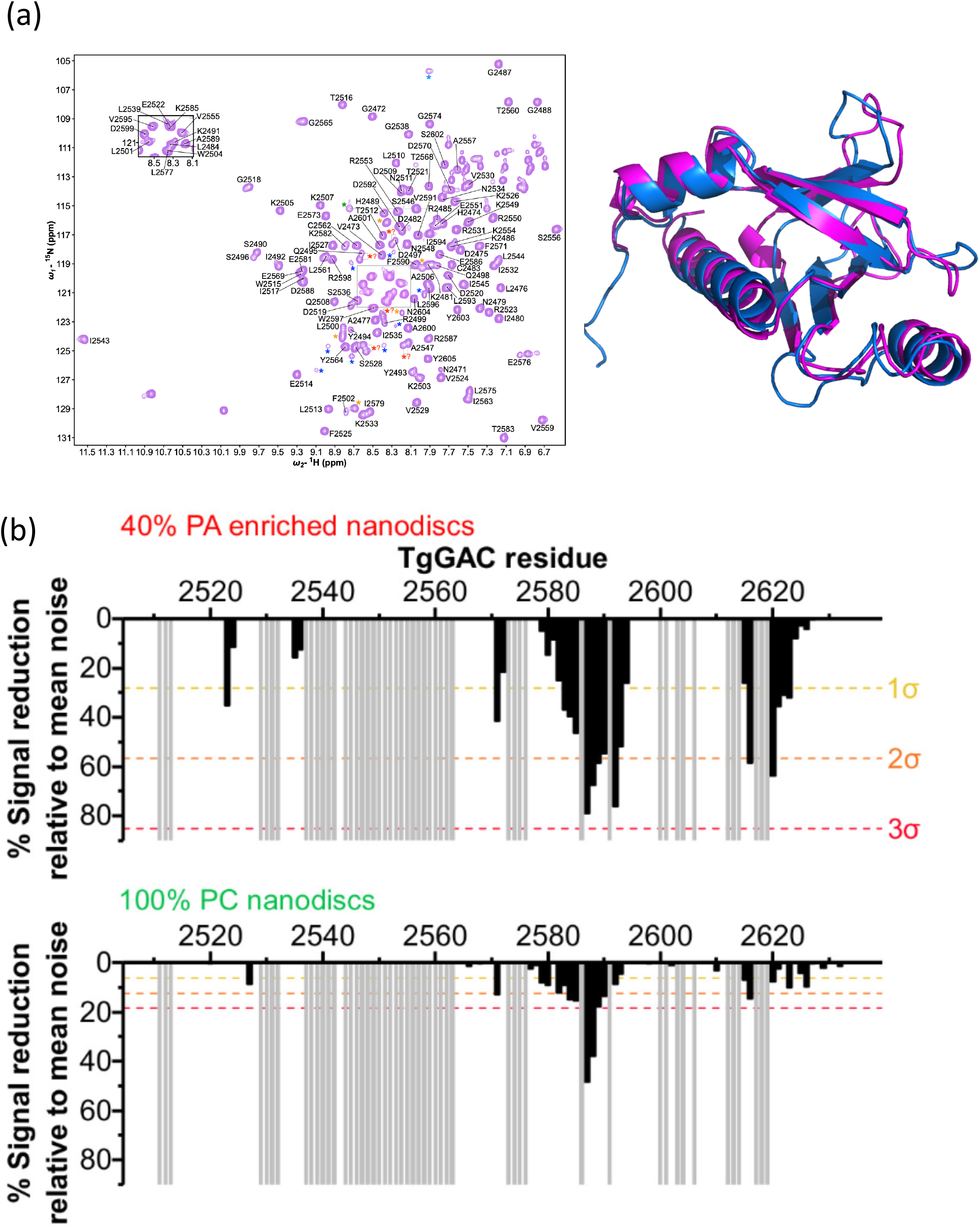
(a) Assigned ^1^H-^15^N HSQC for PfGAC_PH_ and comparison of validated structures for PfGAC_PH_ (magenta) and TgGAC_PH_ (blue) (b) PRE’s with PA-enriched (top) or PC only MSP1D1_H4-5_ nanodiscs

**Figure S5.**
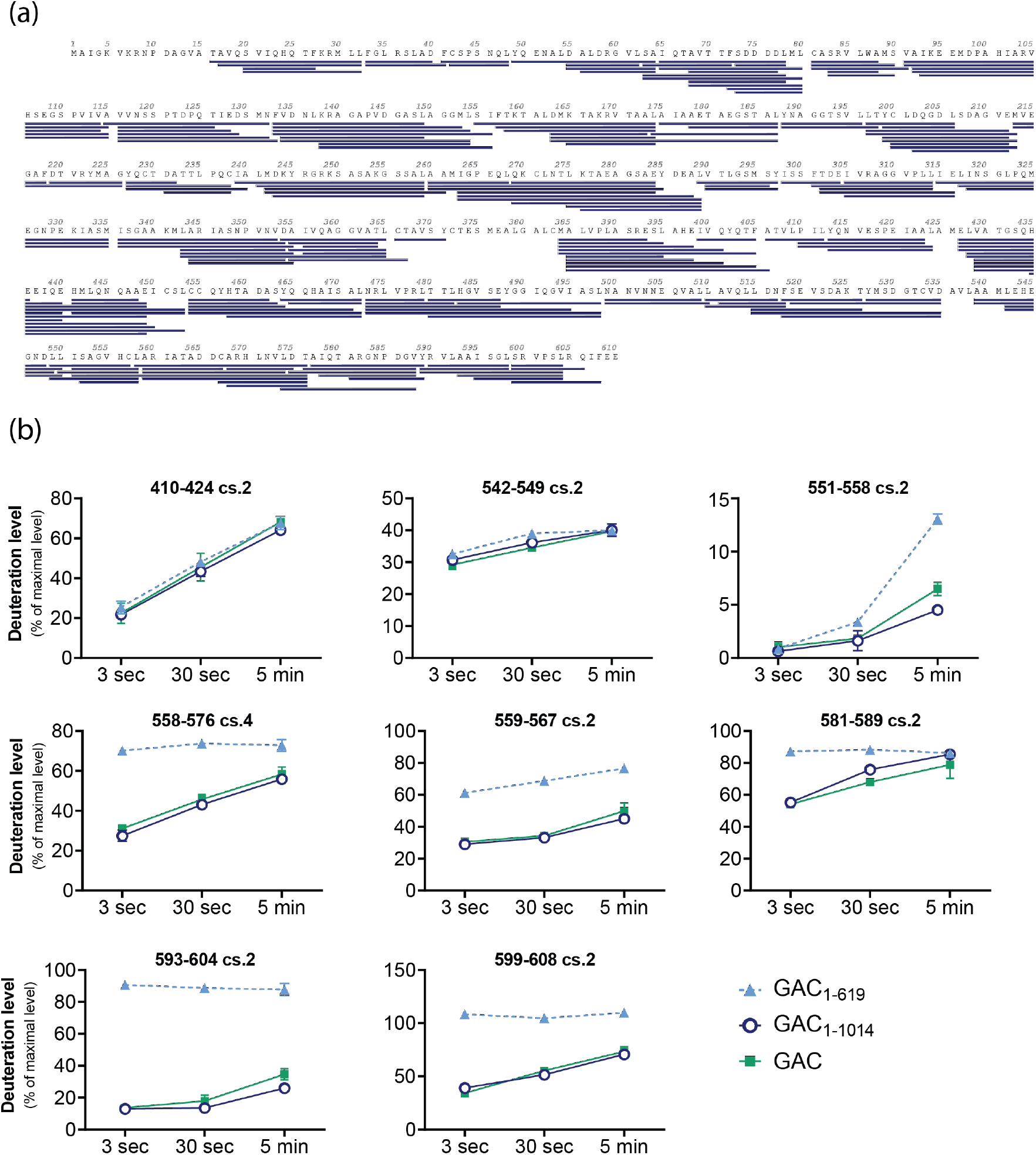
HDX-MS data for GAC_1-619_ construct studied. a) Peptide map showing all peptides analysed in for GAC_1-619_. b) Uptake plots for a selection of peptides

**Table S1.**
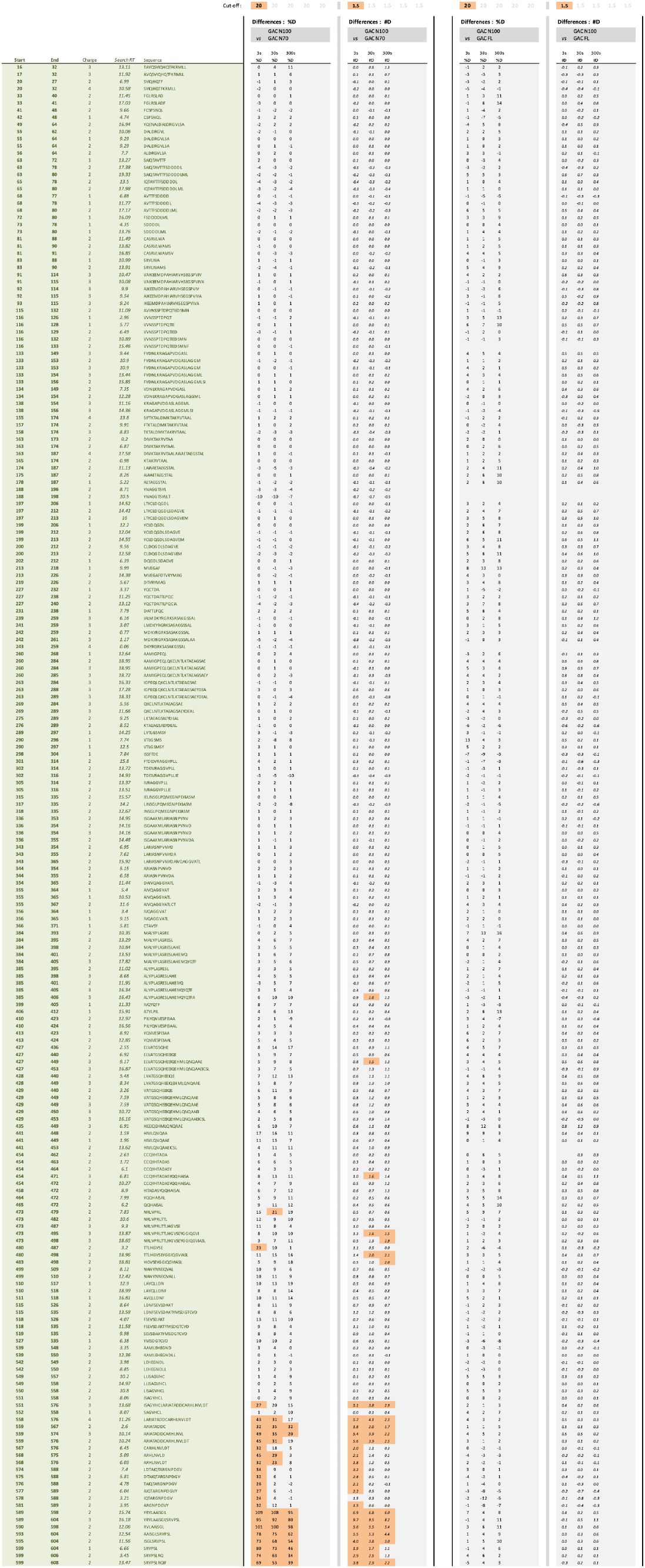
Differences in deuterium uptake between two states. For each state comparison, results are shown for both differences in deuterium uptake by percentage (%D) and by number of deuterons (#D) Largest differences are highlighted in different colors: blue indicates PROTECTION of the peptide and orange indicates increased EXPOSURE (usually associated with allosteric modifications)

**Table S3.**
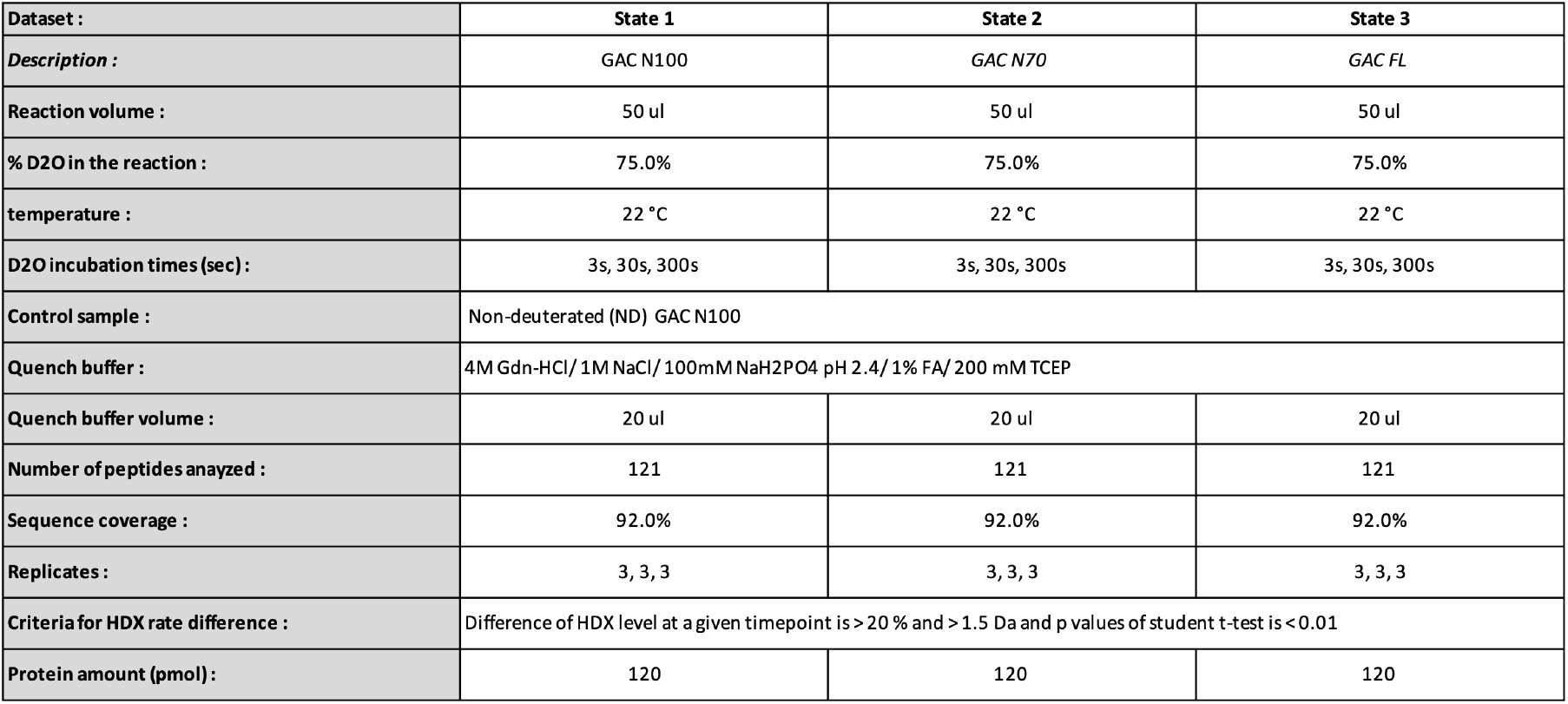
HDX experimental details

**Table S4.**
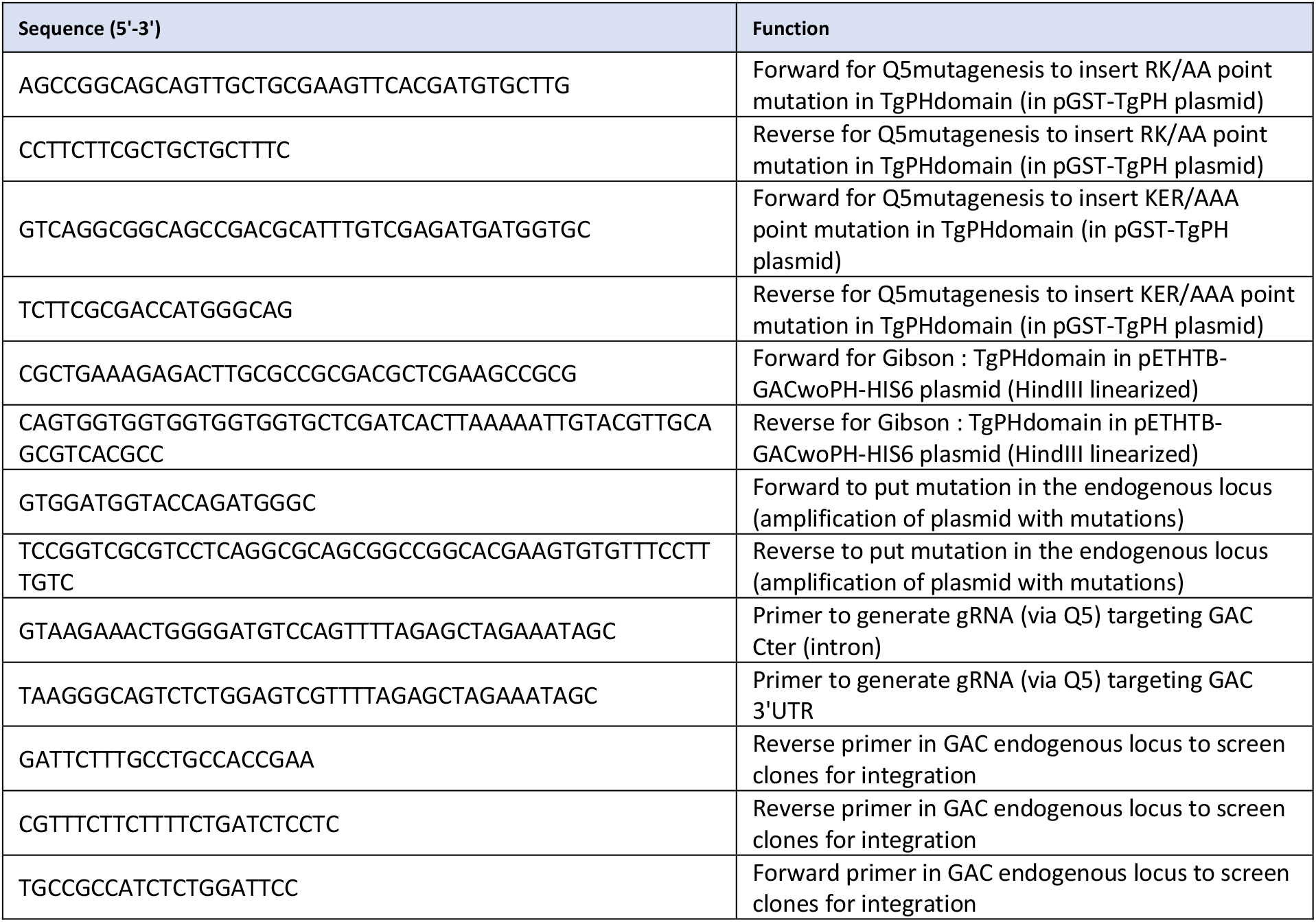
Key DNA primers used generate mutations and the endogenous locus mutants for parasite experiments

## Notes

### Competing Interest Statement

The authors have declared no competing interest.

